# Structure and function of an intermediate GPCR-Gαβγ complex

**DOI:** 10.1101/2024.04.02.587841

**Authors:** Maxine Bi, Xudong Wang, Jinan Wang, Jun Xu, Wenkai Sun, Victor Ayo Adediwura, Yinglong Miao, Yifan Cheng, Libin Ye

**Affiliations:** Department of Biochemistry and Biophysics, University of California, San Francisco, CA 94143; Department of Molecular Biosciences, University of South Florida, 4202 E Fowler Ave, Tampa, FL USA 33620; Department of Pharmacology & Computational Medicinal Program, University of North Carolina at Chapel Hill, 116 Manning Drive, 11004C Mary Ellen Jones Building, Chapel Hill, NC 27599; Department of Molecular and Cellular Physiology, Stanford University School of Medicine, Stanford, CA, USA; Howard Hughes Medical Institute, University of California, San Francisco, CA 94143; H. Lee Moffitt Cancer Center & Research Institute, 12902 USF Magnolia Drive, Tampa, FL, USA 33612

**Author notes:** **Corresponding author** Yinglong Miao, Associate Professor Department of Pharmacology & Computational Medicinal Program, 116 Manning Drive, 11004C Mary Ellen Jones Building, Chapel Hill, NC 27599, **Yifan Cheng, Professor** Department of Biochemistry and Biophysics, University of California, San Francisco, CA 94143; Howard Hughes Medical Institute, University of California, San Francisco, CA 94143, **Libin Ye, Assistant Professor** Department of Molecular Biosciences, University of South Florida, 4202 E Fowler Ave, Tampa, FL, USA 33620; H. Lee Moffitt Cancer Center & Research Institute, 12902 USF Magnolia Drive, Tampa, FL, USA 33612.

## Abstract

Unraveling the signaling roles of intermediate complexes is pivotal for G protein-coupled receptor (GPCR) drug development. Despite hundreds of GPCR-Gαβγ structures, these snapshots primarily capture the fully activated end-state complex. Consequently, a comprehensive understanding of the conformational transitions during GPCR activation and the roles of intermediate GPCR-G protein complexes in signaling remain elusive. Guided by a conformational landscape profiled by ^19^F quantitative NMR (^19^F-qNMR) and Molecular Dynamics (MD) simulations, we resolved the structure of an unliganded GPCR-G protein intermediate complex by blocking its transition to the fully activated end-state complex. More importantly, we presented direct evidence that the intermediate GPCR-Gαsβγ complex initiates a rate-limited nucleotide exchange without progressing to the fully activated end-state complex, thereby bridging a significant gap in our understanding the complexity of GPCR signaling. Understanding the roles of individual conformational states and their complexes in signaling efficacy and bias will help us to design drugs that discriminately target a disease-related conformation.

## Main

G protein-coupled receptors (GPCRs) are the largest family of human membrane proteins, encompassing over 800 distinct members. Due to their vital roles in various (patho)physiological processes, significant efforts have been made to unravel their activation mechanisms, aiming to modulate therapeutic signaling with precision. Recent advancements, especially in single-particle cryo-electron microscopy (cryo-EM), have facilitated the structural determination of hundreds of GPCR-Gαβγ complex since 2017^1–3^. These complex structures primarily represent the receptors in their fully activated end-states. Yet, the GPCR signaling process is more intricate than what these structures alone can depict. Multiple studies have shown that GPCR activation entails transitions through several intermediate complexes, including pre-coupled and partially activated GPCR-G protein complexes^4–12^. Isolating such transient complexes to study their signaling roles presents a significant challenge.

Using the adenosine-A2A receptor (A2AR) as a model system and applying ^19^F quantitative NMR (^19^F-qNMR), we mapped the conformational landscape of this GPCR in response to ligand actions. Within this landscape, we identified two transient active-like intermediates (S3 and S4) that are positioned between the inactive (defined as S1 and S2) and the fully activated states (S5)^13^ (Fig. 1a). Among these conformations, S1 and S2 have been defined by crystal structures ^14,15^, and the S5 state has been characterized by cryo-EM (PDB ID: 6GDG)^16^ (Fig. 1b and Extended Data Fig. 1). The structures of S3 and S4, however, remain uncharacterized. With the R291A point mutation, which we previously determined to trap A2AR uniquely in the intermediate S4 state (Fig. 1)^13^, we have continued to investigate the structure and function of this intermediate state. Our study seeks to answer questions such as whether the intermediate S4 can directly interact with and regulate G protein without transitioning to the S5 state and how the G protein responds to the intermediate S4. Understanding the structural and functional roles of individual intermediate GPCR-G protein complexes will significantly advance our knowledge at the molecular mechanistic level regarding the signaling efficacy, bias, and allostery, which is pivotal for improving drug designs^5,17,18^.

**Fig.1: Identification.**
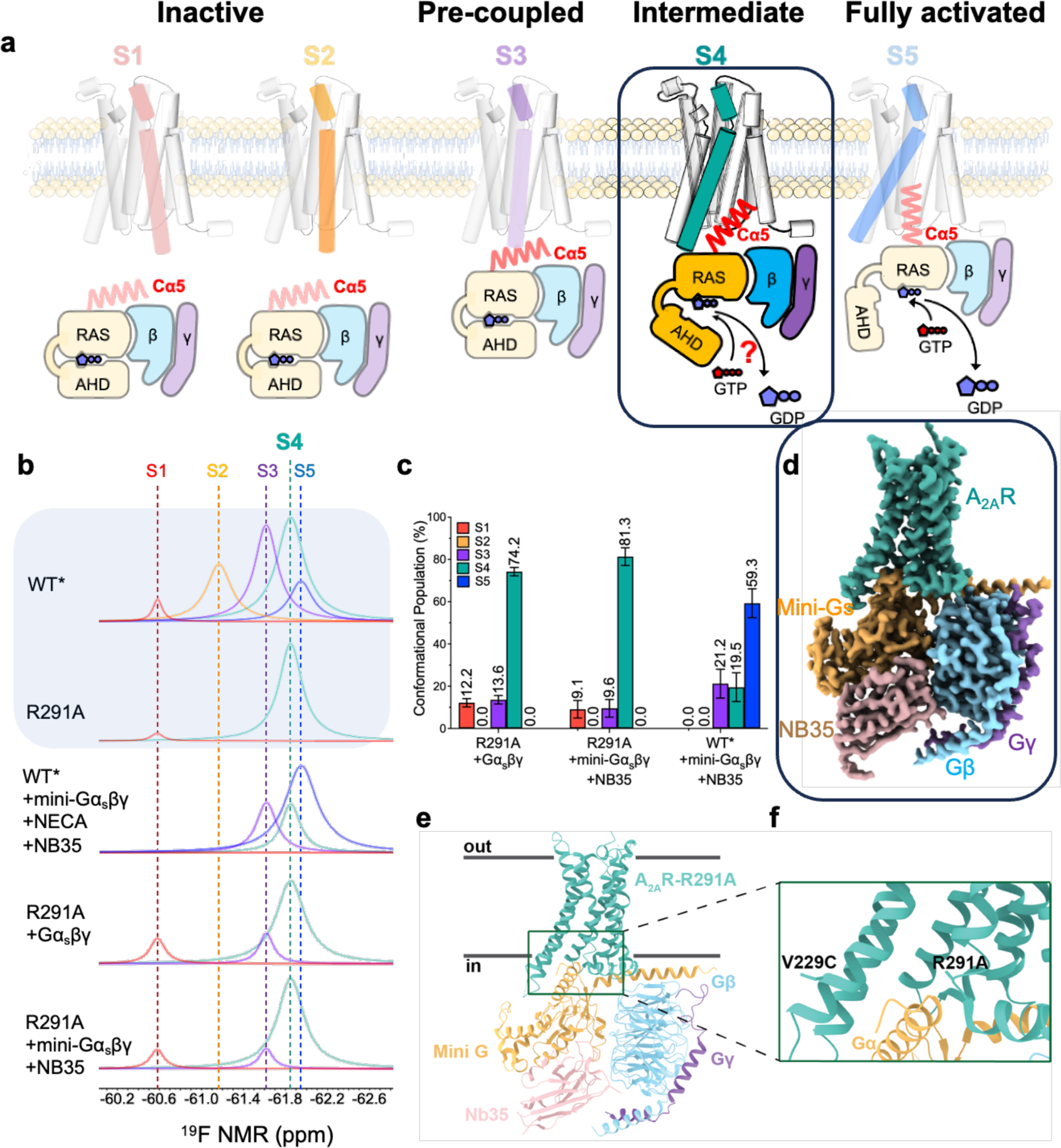
of the intermediate complex investigated in this study. **a,** Proposed activation model of GPCR, including inactive states S1 and S2, pre-coupled complex (S3-Gα_s_βγ), intermediate complex (S4-Gα_s_βγ), and the fully activated complex (S5-Gα_s_βγ), in which S4-Gα_s_βγ was highlighted in teal for this study. **b,** In reference to <u>Extended Data</u> Fig. 1, the deconvoluted conformational profiles probed by^19^F-qNMR for the R291A construct were presented as a function of Gα_s_βγ and mini-Gα_s_βγ. ^19^F-qNMR spectra of WT* and R291A were used as the benchmarks for S1 through S5, adapted from the reference (Wang et al, 2023, Nature Commun). The spectrum of WT*+mini-Gα_s_βγ+NECA+NB35 represents the feature for PDB: 6GDG. **c,** The population distributions of conformational states for R291A+mini-Gα_s_βγ+NB35, R291A+Gα_s_βγ, and WT*+mini-Gα_s_βγ+NB35 (S5-Gα_s_βγ). The SD values were determined based on spectral S/Ns and fitting errors of the deconvolutions. **d,** Ribbon representation of S4-mini-Gα_s_βγ. **e,** Highlighted interfacial section of S4-mini-Gα_s_βγ. **f**, cryo-EM density map of intermediate S4-mini-Gα_s_βγ complex.

### S4 state binds to Gαsβγ and initiates nucleotide exchange

We first examined whether the WT*-R291A mutant, which is trapped in the S4 state, can bind to the Gαβγ. Here, the S4 trapped mutant was introduced to a previously defined construct (A2AR-316-V229C, denoted as WT*),^13^ and we referred to WT*-R291A simply as R291A. We incubated ^19^F-labeled R291A with Gαβγ, subjected it to ^19^F-NMR, and measured the linewidth of the NMR resonance. Indeed, we observed a linewidth broadening for the S4 resonance compared to the receptor alone, suggesting direct binding of Gαβγ to A2AR in the S4 state (Extended Data Fig. 2a). This was further confirmed by a complex band of apo R291A-Gαsβγ in native-PAGE (Extended Data Fig. 2b).

Next, we assessed how the S4 regulates G protein function by performing nucleotide hydrolysis, binding, and exchange assays. As shown in Fig. 2a, the R291A mutant-mediated GTP hydrolysis from the G protein occurred at a much slower rate compared to the full agonist-bound WT*-Gαsβγ (representing the S5-Gαsβγ). However, the addition of full, partial, or inverse agonists did not alter the GTP hydrolysis level substantially in the R291A mutant. This ligand-independent hydrolysis behavior is in stark contrast to the wild-type construct, where ligand binding dramatically changes the hydrolysis level^19,20^. To further investigate the reduced hydrolysis level, we measured the initial rates and Michaelis constants of GTP hydrolysis. The initial rate of GTP hydrolysis mediated by R291A was only one-twentieth of that mediated by WT* (Fig. 2b), while the Michaelis constant exhibited a similar pattern (Fig. 2c).

**Fig. 2:**
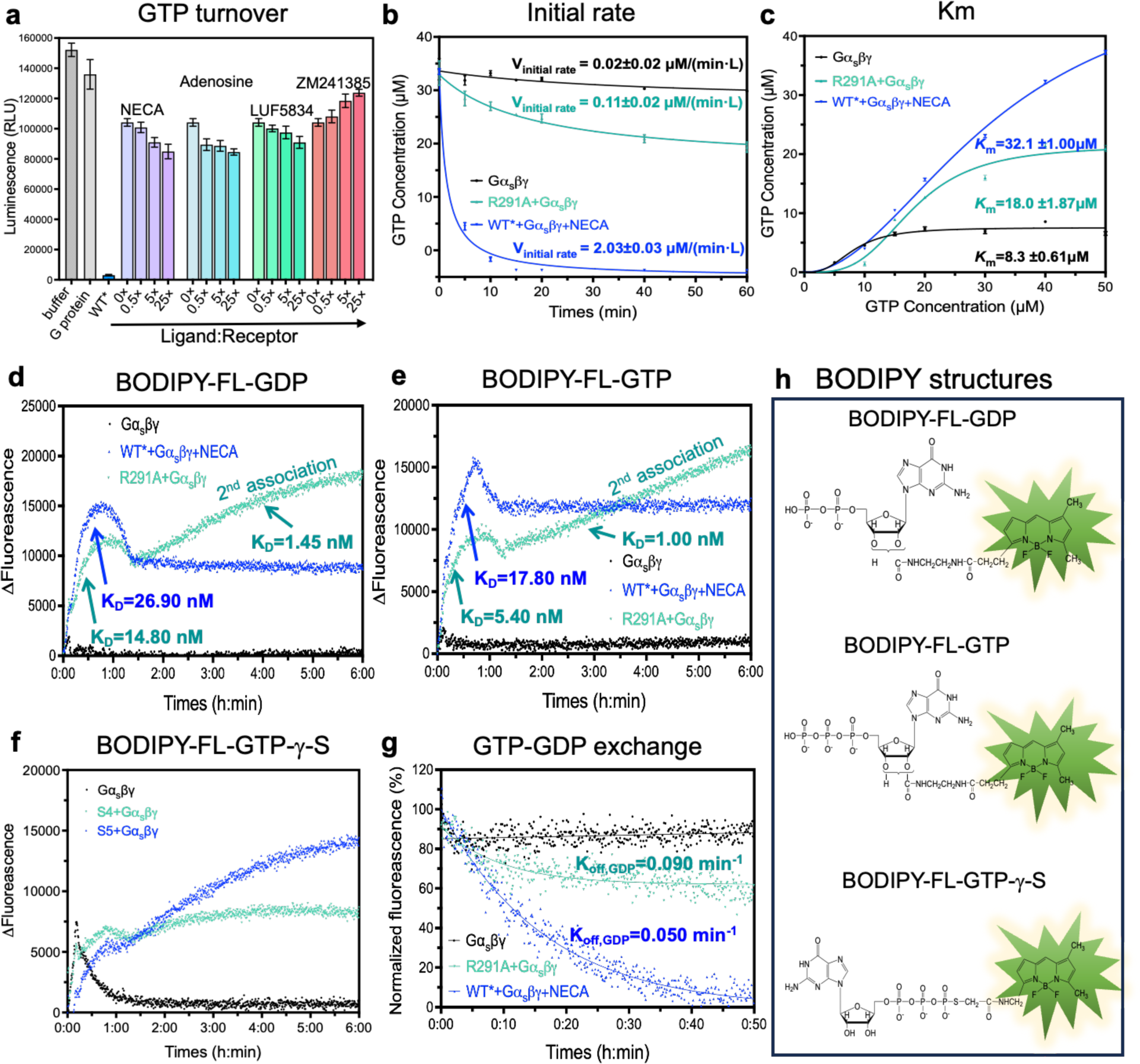
S4 mediated GTP hydrolysis and nucleotide exchange. **a,** GTP hydrolysis of the S4 mediated Gα_s_βγ as a function of inverse, partial, and full agonists, in reference to Gα_s_βγ alone (negative control) and the S5 mediated Gα_s_βγ (positive control). **b,** Time course of GTP hydrolysis of the S4 mediated Gα_s_βγ, in reference to Gα_s_βγ alone (negative control) and the S5 mediated Gα_s_βγ (positive control); the initial rate of each catalysis was also calculated as presented. **c,** Km values for S4 mediated Gα_s_βγ, in reference to Gα_s_βγ alone (negative control) and the S5 mediated Gα_s_βγ (positive control). **d,** BODIPY-FL-GDP binding assay. **e,** BODIPY-FL-GTP binding assay. **f**, BODIPY-FL-GTP-γ-S binding assay **g,** GTP-GDP exchange rate comparison. **g**, The structures of BODIPY-FL-GDP, BODIPY-FL-GTP, and BODIPY-FL-GTP-γ-S.

We proposed a simple model to explain the limited GTP hydrolyzing capacity of the S4-mediated G protein, where nucleotide binding to G protein is a two-step process. A nucleotide GTP initially binds to a site with low affinity and hydrolysis (site1) in the intermediate S4-G protein complex. Upon the transitioning from S4 to S5, the G protein changes its conformation to allow the GTP to access the site 2, the canonical nucleotide-binding site. We tested this model by examining the binding of BODIPY-FL-GDP (Figs. 2d and 2h). The BODIPY-FL-GDP bound to the WT*-mediated G protein exhibited a higher KD than that of the R291A-mediated G protein, although both reached equilibrium at 1.5 hrs. For the R291A-mediated G protein, a secondary association event was observed after 1.5 hrs with a KD value 10 times lower, which was not the case observed in the WT*. We attribute this difference to the restricted accessibility of the site 2, caused by spatial constraints resulting from the partially open AHD domain in the S4 mediated G-protein, while this site is freely accessible in the WT*. A similar pattern was noted in BODIPY-FL-GTP binding assays, where BODIPY-FL-GTP showed more efficient binding to the S5-mediated G protein with a higher KD value (Fig. 2e and 2h). The spatial limited association process was also evident in BODIPY-FL-GTP-γ-S (Fig.2f) in both S5 and S4 mediated G protein because the BODIPY head was linked to γ-phosphate, which is meant to be inserted into the site 2. The bulky BODIPY head insertion restricted exchange and hydrolysis. Titration of GTP into the BODIPY-FL-GDP equilibrium system led us to conclude that the S4-mediated G protein has a reduced nucleotide exchange rate compared to the S5-mediated G protein (Fig. 2g). A previous study of β2AR indicated that GDP release occurs much faster than the formation of fully activated complex^21^, while MD simulations suggested that GDP is released when the Gαs is not fully opened^22^. Collectively, our data suggests the intermediate S4 state mediated G-protein initiates nucleotide exchange before transitioning to the fully activated S5 state, albeit at a reduced rate.

### Structure of the ligand-free intermediate S4-mini-Gαsβγ complex

Building upon the two-step nucleotide binding model, we hypothesized that the S4 state not only adopts a conformation distinct from S5 but also binds to the G protein in a manner conformationally different from the S5-bound state. To test this hypothesis and enhance our understanding of the molecular mechanisms involved, we aimed to determine the structure of the intermediate S4 state in complex with the G protein.

To capture the structure of intermediate S4 state in complex with G proteins, we assembled a stable complex by incubating the R291A mutation with mini-Gαs, known as mini-Gαs399, and the Gαs-stabilizing nanobody, NB35, in conjunction with Gβ1γ2. As evidenced by ^19^F-qNMR, the receptor preserves the conformation of the S4 state in complex with G proteins (Fig.1b and 1c). We determined a cryo-EM structure of this S4-mini-Gαsβγ complex at a 2.8 Å resolution, revealing the key interaction sites between A2AR and mini-Gαsβγ without ligand (Figs. 1d-f and Fig.3a). For a comprehensive understanding of our methods, we have detailed the sample preparation, cryo-EM data acquisition, and processing steps in Extended Data Figs. 3-6, and Supplementary Table 1.

Comparisons of the S4-mini-Gαsβγ complex with the S5-mini-Gαsβγ complex (PDB ID: 6GDG), as depicted in Supplementary videos 1 and 2, reveal key differences between the two states. The extracellular orthosteric binding pocket of the intermediate complex S4-mini-Gαsβγ is less compact than of the S5-mini-Gαsβγ (Fig. 3b), yet more constricted compared to the inactive state^14,15^. Transitioning from the inactive to the intermediate state S4 involves significant inward movements of all domains relative to TM1 in the extracellular domains. The shift from the intermediate to the fully activated complex results in observable movements of these domains. This is consistent with the ^19^F-qNMR study, which recorded a substantial chemical shift (∼400 Hz) when the receptor transitioned from the inactive states S1-2 to intermediate S4, while the shift from S4 to S5 resulted in a minor chemical shift (∼60 Hz) (Fig.3f). Closer inspection of the TM6 domain, using the ^19^F-tag labeling site as a reference, revealed a clockwise rotation consistent with the ^19^F-qNMR resonance, where the NMR signal for the S4 state is at a lower field than the S5 state. Indeed, the conformation of the intermediate S4 state is thus on the trajectory from the inactive S1-2 to the fully activated S5 states.

**Fig. 3:**
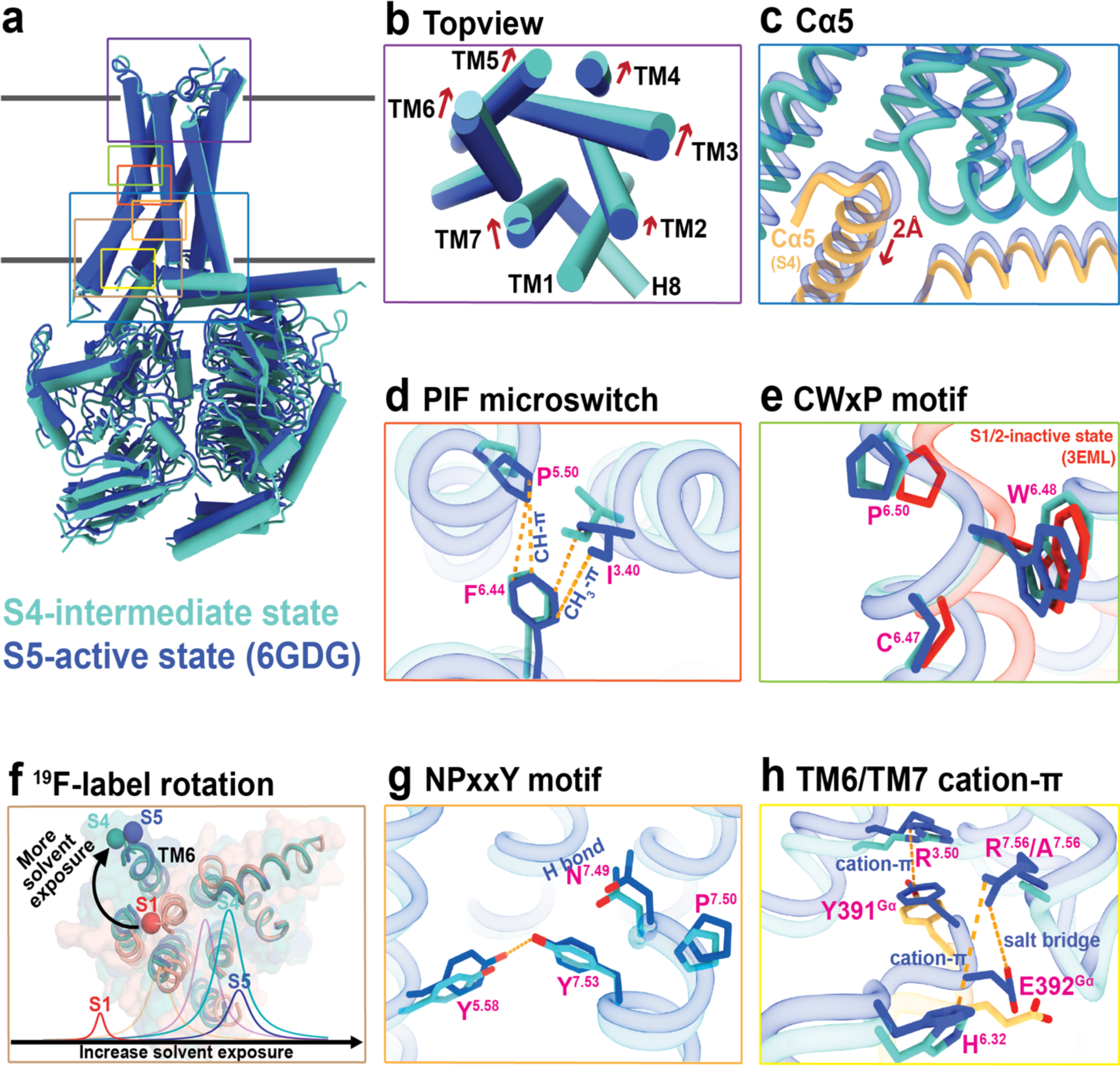
Structural features of the receptor in intermediate complex. **a,** Superimposed cylinder representations of S4-mini-Gα_s_βγ and S5-mini-Gα_s_βγ. **b,** Top view of ligand binding domain of the S4 state compared to the S5 state. **c**, Cα5 motion of Gα_s_. **d**, PIF microswitch. **e**, CWxP motif. **f**, ^19^F-tag rotation, and corresponding NMR spectrum. **g,** NPxxY motif. **h**, TM6 and TM7 cation-π interaction in two complexes.

One of the key microswitches in the receptor is the <u>P</u>^5^<u>^.50^I</u>^3^<u>^.40^F</u>^6.44^ motif, which is a crucial hydrophobic switch between TM5, TM3 and TM6 and is implicated in receptor activation^1,23^. In the fully activated receptor, P^5.50^ and F^6.44^ are in cis positions, forming a strong CH-π interaction^24^, and the CH3 group of I^3.40^ also forms a robust CH3-π interaction with F^6.44^, stabilizing the fully activated conformation. In contrast, these interactions are weakened in the S4-mini-Gαsβγ complex because P^5.50^ and I^3.40^ are oriented away from F^6.44^ (Fig. 3d). Another key microswitch is the <u>CW</u>^6.48^<u>xP</u> motif, where P^6.50^ acts as a hinge in TM6 and facilitates the opening of the cytoplasmic cavity during activation^25^. Our cryo-EM structure clearly shows P^6.50^ in an “intermediate” position, bridging the inactive and active states (Fig. 3e). The third microswitch, the <u>NPxxY</u>^7.53^ on TM7, along with a conserved tyrosine Y^5.58^ in TM5, stabilizes the receptor’s active state^26–28^. As illustrated in Fig. 3g, a strong H-bond is observed between Y^5.58^ and Y^7.53^ in the S5-mini-Gαβγ complex. This interaction draws Y^5.58^ on TM5 closer to TM7, maintaining the compactness of the TM bundle in the S5-G protein complex, as shown in the animation from the S4-mini-Gαsβγ transitioning to the S5-mini-Gαβγ (Supplementary videos 1 and 2). Therefore, transitioning from the S4- to the S5-mini-Gαsβγ strengthens the H-bond interaction as Y^5.58^ moves closer to the TM7 domain, aiding in the insertion of the G protein into the cavity. Our analysis indeed supports the idea that the S4 conformation can regulate G protein signaling to some degree without transitioning to the fully activated S5 state.

Next, we examined the interfacial interaction between Cα5 helix of mini-Gαs (Fig. 3c) and the transmembrane bundle comprising TM3, TM6 and TM7 domains - critical components in regulating G protein nucleotide exchange. As highlighted in Fig. 3c, the S4-mini-Gαsβγ adopts a slightly more open conformation than the S5-mini-Gαsβγ. The Cα5 helix is significantly rotated outward and retracted from the G protein-binding pocket, accompanied by the disengagement of the αN helix of the Gαs protein from the H8 helix of the receptor. This retraction measures approximately 2 Å and the rotation is at a 2° spinal clockwise angle from the S5-mini-Gαsβγ. This motion is further detailed in Fig. 3h, which shows the primary interactions at the junction composed of TM3, TM6, TM7 domains, and the Cα5 segment from mini-Gαs. During the activation process, the clockwise rotation of TM6 and counterclockwise rotation of Cα5 culminate in a compact end-state complex. The transition from the S4- to the S5-mini-Gαsβγ entails a further insertion of the G protein and an inward rotation of the receptor domain, evidenced by the formation of the salt bridge R^7.56^-E392^Gα^ and the cation-ρε interaction between R^7.56^ and H^6.32^, coupled with a strengthened cation-ρε interaction between R^3.50^ and Y391^Gα^. However, the R^7.56^ to A^7.56^ mutation eliminates this interaction.

### Molecular basis of A2AR function in the S4 state

To validate the conformational transition between the S4 and S5 states, we applied all-atom Gaussian accelerated MD (GaMD) simulations, starting from atomic structure of the S5-mini-Gαsβγ complex (6GDG) with a R291A mutation modeled in silico. Indeed, the R291A mutation facilitated transition of the 6GDG structure (S5 state, Fig.1b) to an energy minimum conformation that closely matches our new cryo-EM structure. We refer to this as cS4 state, where “c” stands for computational model, and here the Cα5 helix in Gαs retracted from the receptor by ∼2 Å. The center-of-mass distance between the receptor NPxxY motif and the last five residues of Cα5 helix increased from ∼13.2 Å in the “S5” state to ∼15.3 Å in the “cS4” state (Fig. 4a and Extended Data Fig. 7). As a control, GaMD simulations showed that the cWT-mini-Gαsβγ complex remained stable, mainly sampling one low-energy state, corresponding to the S5 (Fig.4b). These simulations independently validated the conformation of the intermediate S4 state determined from single particle cryo-EM.

**Fig. 4:**
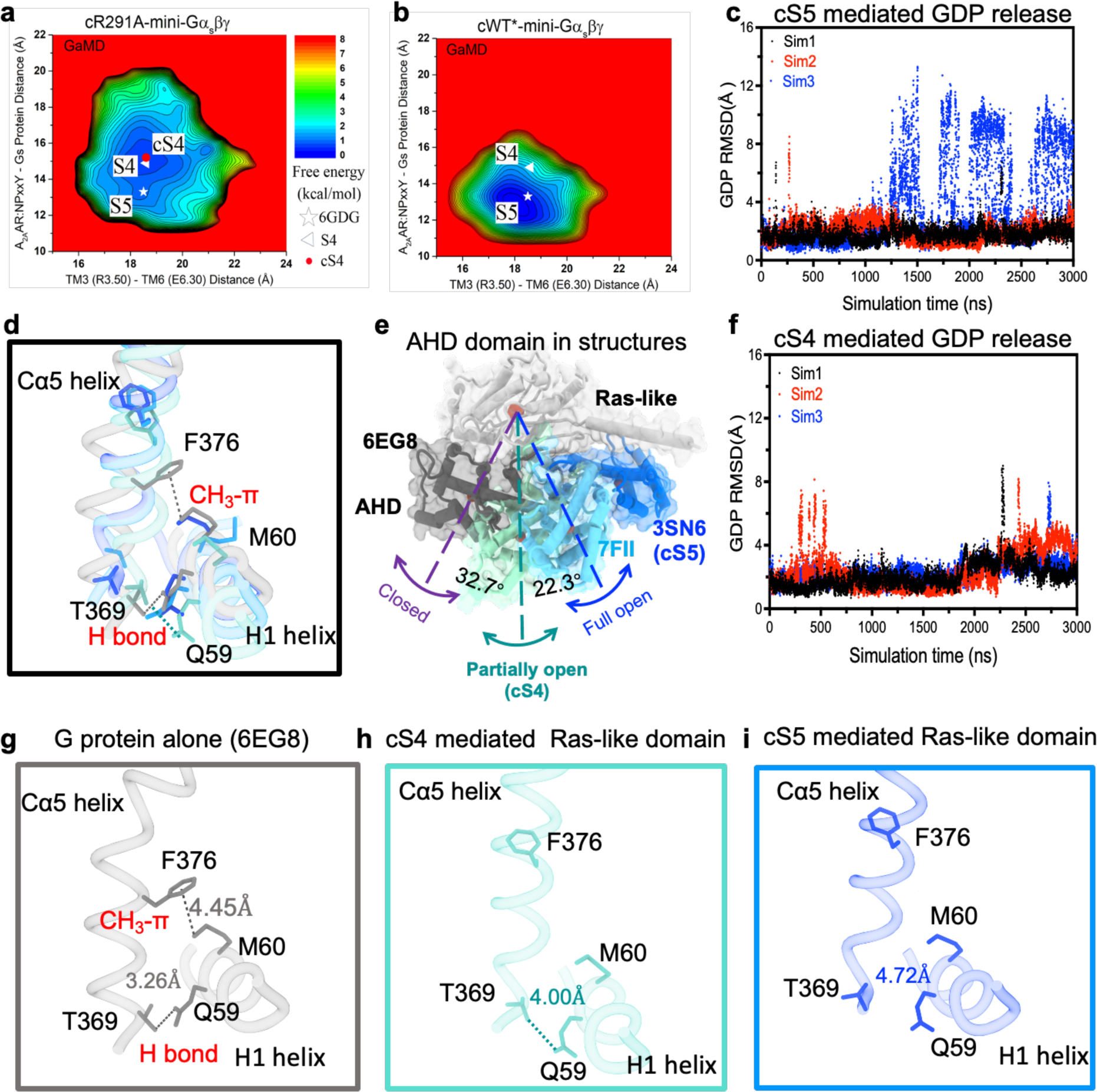
Structural features of A_2A_R and G protein in GaMD simulation. **a,** 2D free energy profiles of the cR291A-mini-Gα_s_βγ complex system. **b**, 2D free energy profiles of the cWT*-mini-Gα_s_βγ complex systems. The white triangle and asterisk indicate cryo-EM structures of the S5 and S4 states, respectively. **c**, Root-mean square deviation (RMSD) of GDP relative to the initial position in the GaMD simulation for cS5 mediated G protein. **d**, Superimposed positions of Cα5 and H1 helices when G protein is inactive, partial activated, and fully activated. **e**, The positions of AHD domain when G protein is inactive, partial activated, and fully activated. **f**, RMSD of GDP relative to the initial position in the GaMD simulation for cS4 mediated G protein. **g**-**i**, The relative positions of Cα5 and H1 helices when G protein is inactive (6EG8), partial activated (cS4 mediated), and fully activated (cS5 mediated).

Furthermore, we performed additional GaMD simulations to examine GDP release. In the cWT-Gαsβγ system, GDP was observed moving away from the initial binding site by up to ∼12 Å repeatedly (Fig.4c). In contrast, GDP underwent significantly smaller movements up to only ∼8 Å in the cR291A-Gαsβγ system (Fig. 4f). Free energy calculations also indicated a “GDP Released” state in the cWT-Gαsβγ system and only a “Partially Released” state in the cR291A-Gαsβγ system (Extended Data Figs. 8a-c), where the AHD domain of Gαs transitioned from the “Open” to “Partially Open” conformation, with the orientation angle between the AHD and Ras-like domain decreasing to ∼30° (Extended Data Figs. 8d-e) ^1,29^. Together, these observations suggest that the S4 state mediated G protein has a reduced capacity for GDP release compared to the S5 state, aligning with the nucleotide exchange data shown above.

### A limited nucleotide exchange model for intermediate GPCR-Gαsβγ complex

A mechanistic model is proposed to understand the rate-limited nucleotide exchange in the intermediate GPCR-G protein complex (Fig.5). The transition from the intermediate to the fully activated complex involves a conformational change at the interface, where the TM6 helix of the receptor rotates clockwise by 8° while the Cα5 helix from the Ras-like domain rotates anticlockwise by 2°. This leads to a more compact interaction between the receptor and G protein, coupled with a 2 Å uplift of the Cα5 helix. This uplift results in the disengagement of the H1 from the Cα5 helix, facilitating the separation of AHD domain from Ras-like domain and thereby exposing the bound GDP at site 2. Simultaneously, this process assists the relocation of GTP from site 1 to site 2, replacing GDP. However, in the S4-mediated G protein, the substitution of R^7.56^ with alanine (A^7.56^) results in the loss of both the intramolecular cation-π (R^7.56^-H^6.32^) interaction and the intermolecular salt bridge (R^7.56^-E392^Gα^). Instead, a partially opened AHD domain is seen (Fig.4e), which can be attributed to a weak electrostatic H-bond (Q59-T369, 4 Å)^30^ between Cα5 and H1 helices. While in the closed state of the AHD domain, an additional CH3-π interaction between F376 and M60 plays a significant role (Fig.4d and Figs.4g-i). Of note, in all complex structure resolved so far, the crystallographic complex structure of β2AR-Gαβγ represents the sole fully activated complex structure, in which the AHD domain with an open of 88° while all others are between 55-65°. This could be the effect from crystallographic packing in the crystallization process ^31^. Collectively, these findings provide a molecular basis for the limited nucleotide exchange observed in the intermediate GPCR-G protein complex.

**Figure 5:**
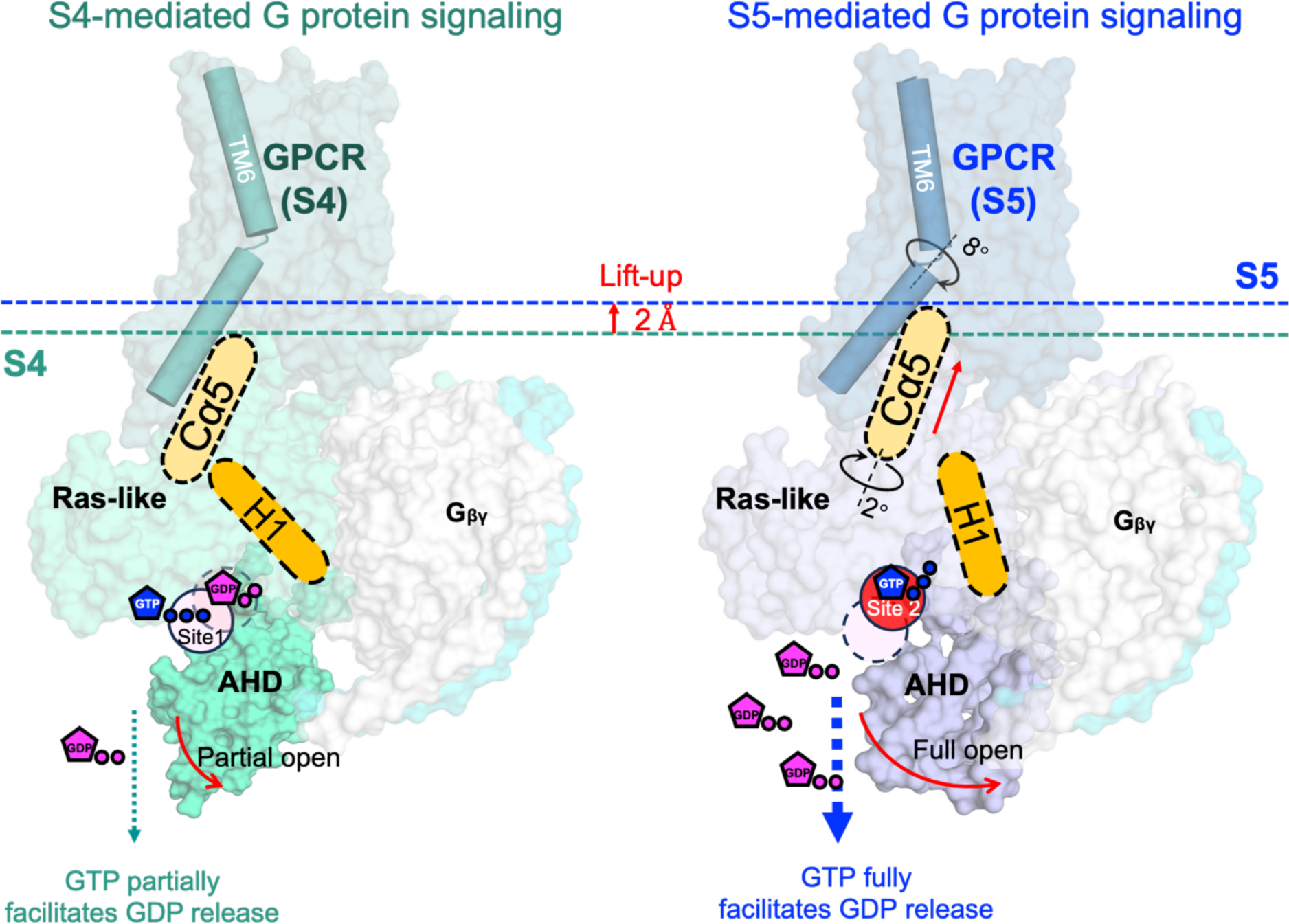
A limited nucleotide exchange model for intermediate GPCR-Gαsβγ complex. In the S5-mediated Gα the interaction between Cα5 and H1 helices was interrupted while this interaction is partially maintained through a weakened H-bond between Cα5 and H1 helices in Ras-like domain.

## Discussion

Previous MD simulation has indicated that GTPγS and GDP exhibited different contact patterns with G protein^32^. A recent study also demonstrated a transition process of GTP nucleotide in a β2AR-regulated G protein from a “loose” interaction site to the final nucleotide-binding pocket by 2 Å^33^. Our study provides evidence for a 2 Å distance dislocation of nucleotide, regulated by a “lift” of the Cα5 tail by 2 Å, which is essential for dislodging the AHD domain of the G protein from the Ras-like domain. This facilitates the opening of the canonical site 2, accommodating GTP for free-nucleotide exchange and hydrolysis^34^. However, the precise location of site 1 en route from the S4-Gαβγ to the S5-Gαβγ in the GPCR activation process and its specific functional role in G protein signaling remain unclear and require further characterization in the future.

GPCR activation is known to involve the transition through multi-states^4–12^. However, transient intermediate states represent higher energy substates from energy landscape perspective and are thus challenging to characterize, especially with regard to obtaining transient GPCR-G protein complexes and studying their functions. Our current study demonstrates a strategy whereby intermediate conformational states can be identified by ^19^F-qNMR, trapped by point mutation, and structurally and functionally characterized in their complexes with G proteins. Although we used A2aR as a model system, it is clear that such strategy of utilizing 19F-qNMR-guided cryo-EM, in which 19F-NMR serves as conformational indicator, can be broadly applied to study transient complexes of other GPCRs and proteins of interest during activation or inhibition processes. Deciphering the functions of intermediate states and their complexes enables a comprehensive understanding of GPCR signaling complexity and the exploration of alternative therapeutic approaches by targeting a specific disease-related conformation(complex). As the advance of our knowledge on structures and functions of individual conformational states and their complexes, it will become possible to design drug based on their conformational selectivity on the receptor, considering signaling bias originated from conformational bias ^5^.

## Methods

### Plasmid construction and transformation

The full-length human A2AR gene, originating from construct pPIC9K_ADORA2A, was generously provided by Prof. Takuya Kobayashi (Kyoto University, Kyoto, Japan). The C-terminally truncated construct A2AR_316, constructed in our previous study, has an integrated FLAG tag on the N-terminus and a poly-his tag on the C-terminus^7^. Based on this construct, the mutations V229C and R291A were described elsewhere^13^. All constructs were sequenced by a facility at Eurofins genomics, with the AOX1 primer pair of PFAOX1 and PRAOX1. Freshly prepared competent cells of strain *Pichia Pastoris* SMD 1163 (*Δhis4 Δpep4 Δprb1,* Invitrogen) were electro-transformed with *Pme*I-HF (New England Biolabs) linearized plasmids containing different mutant genes using a Gene Pulser II (Bio-Rad). High-copy clone selection was performed using an in-house protocol described previously^8,35^. A high-yield construct was then screened by an immunoblotting assay with both anti-FLAG and anti-Poly-his for further large-scale expression screening.

### Receptor expression, purification, and labeling

The screened WT* and mutants R291A, were pre-cultured on YPD [1% (w/v) yeast extract, 2% (w/v) peptone and 2% (w/v) glucose] plates containing 0.1 mg/mL G418. A single colony for each construct was inoculated into 4 mL YPD medium and cultured at 30 °C for 12 hours, then transferred into 200 mL BMGY medium [1% (w/v) yeast extract, 2% (w/v) peptone, 1.34% (w/v) YNB (yeast nitrogen base) without amino acids, 0.00004% (w/v) biotin, 1% (w/v) glycerol, 0.1 M PB (phosphate buffer) at pH 6.5] and cultured at 30 °C for another 30 h. The cells were then transferred into 1 L of BMMY medium [1% (w/v) yeast extract, 2% (w/v) peptone, 1.34% (w/v) YNB without amino acids, 0.00004% (w/v) biotin, 0.5% (w/v) methanol, 0.1 M phosphate buffer at pH 6.5, 0.04% (w/v) histidine and 3% (v/v) DMSO, 10 mM theophylline] at 20 °C. 0.5% (v/v) methanol was added every 12 h. 60 h after induction by methanol, cells were harvested for purification.

The cell pellets were collected by centrifugation at 4,000 ×g for 20 minutes and washed one time with washing buffer (50 mM HEPES, 10% glycerol, pH 7.4) before the addition of breaking buffer (50 mM HEPES, pH 7.4, 100 mM NaCl, 2.5 mM EDTA, 10% glycerol) in a ratio of 4:1 (buffer: cells). The resuspended cell pellets were subject to disruption 3 times using a Microfluidizer at a pressure of 20,000 psi. Intact cells and cell debris were separated by low-speed centrifugation (8,000 ×g) for 30 minutes. The supernatant was collected and centrifuged at 100,000 ×g for 2 h, and the precipitated cell membrane was then immediately dissolved in membrane lysis buffer (50 mM HEPES, pH 7.4, 100 mM NaCl, 0.5% LMNG-3 (Lauryl Maltose Neopentyl Glycol) and 0.1% CHS (cholesteryl hemisuccinate)) with rotation 2 h or overnight at 4 °C until the membrane was dissolved. Subsequently, Talon resin (Clontech) was added to the solubilized membranes and incubated for at least 2 h or overnight under gentle agitation.

The A2AR-bound Talon resin was washed twice with a buffer of 50 mM HEPES, pH 7.4, 100 mM NaCl, 0.02% LMNG-3 and 0.002% CHS and resuspended in the same buffer. The A2AR-bound Talon resin was then resuspended in buffer made of 50 mM HEPES, pH 7.4, 100 mM NaCl, 0.02% LMNG-3 and 0.002% CHS, and combined with 10-20 fold excess of the NMR label (2-bromo-*N*-(4-(trifluoromethyl)phenyl)acetamide, BTFMA, Apollo Scientific, Stockport, UK) under gentle agitation overnight at 4 °C. Another aliquot of NMR label was then added and incubated for an additional 6 h to ensure complete labeling. The A2AR-bound Talon resin was washed in a disposable column extensively with buffer containing 50 mM HEPES, pH 7.4, 100 mM NaCl, 0.02% LMNG-3 and 0.01% CHS, and apo A2AR was then eluted from the Talon resin with 50 mM HEPES, pH 7.4, 100 mM NaCl, 0.02% LMNG-3 and 0.01% CHS, 250 mM imidazole and concentrated to a volume of 5 mL. The XAC-agarose gel and A2AR were then incubated together for 2 h under gentle agitation. The functional A2AR was eluted with 50 mM HEPES, pH 7.4, 0.02% LMNG-3, 0.002% CHS, 100 mM NaCl, 20 mM theophylline. The eluted samples were concentrated to 1 mL by centrifugal filtration (MWCO, 3.5 KDa), and an extensive dialysis was performed to remove the XAC in the sample. The functional apo A2AR was then prepared for NMR. All receptors described in this manuscript were purified using poly-his resin followed with a ligand-column, in which the A2AR antagonist xanthine amine congener (XAC) was conjugated to Affi-Gel 10 activated affinity media.

### Preparation of mini-Gαsβγ heterotrimer

The plasmid for mini-Gαs399 was generously provided by Drs. Christopher G. Tate from MRC Laboratory of Molecular Biology, Cambridge and Javier García Nafría from Institute for Biocomputation and Physics of Complex Systems (BIFI) and Laboratorio de Microscopías Avanzadas (LMA), University of Zaragoza. It was expressed in the *E. coli* strain BL21(DE3). Cells were collected by centrifugation at 4,000 ×g for 20 min and lysed by sonication. After another centrifugation, the supernatant was purified by Talon resin. The sample was loaded into a HiLoad16/60 column to obtain purified mini-Gαs protein. The purified protein was concentrated to 3 mg/mL and flash frozen in liquid nitrogen and stored at -80 °C for further use. The expression and purification of the respective components and assembly to make the complex containing mini-G**α**sβ1γ2, and the preparation of nanobody Nb35, were all performed following the protocols described previously ^1,36,37^.

### 19F NMR experiments

NMR samples typically consisted of 280-300 μL volumes with 20-50 μM A2AR in 50 mM HEPES buffer and 100 mM NaCl, doped with 10% D2O. The receptor was stabilized in 0.02% LMNG-3 and 0.002% CHS. All ^19^F NMR experiments were performed on a 600 MHz Varian Inova spectrometer using a ^19^F dedicated resonance probe. Typical experimental setup included a 16 μs 90° excitation pulse, an acquisition time of 200 ms, a spectral width of 15 kHz, and a repetition time of 1 s. Most spectra were acquired with 15,000-50,000 scans. Processing typically involved zero filling, and exponential apodization equivalent to 15 Hz line broadening.

### GTPase hydrolysis assay

The GTPase hydrolysis assay was analyzed using a modified protocol of the GTPase-Glo^TM^ assay (Promega)^38^. The reaction was started by mixing 300 nM Gαsβγ with the purified receptors in varied concentrations with a final volume of 10 μL in the buffer containing 50 mM HEPES, pH 7.4, 100 mM NaCl, 0.002% CHS, 0.02% LMNG-3. For the GTP hydrolysis capacity of the S4 state as a function of ligand measurement, 5x and 25x ligand compared to receptor concentration was added. After 30 min incubation at room temperature, 10 μL 2xGTP-GAP solution containing 10 μM GTP, 1 mM DTT and the cognate GAP was added to each well, followed with a 120 min incubation at room temperature. For the Michaelis–Menten constant measurement, the 2xGTP-GAP solution containing 5-50 μM GTP was used. Then, 20 μL reconstituted GTPase-Glo^TM^ reagent containing 5 μM ADP was added to each sample and incubated for another 30 min at room temperature with shaking. Luminescence was measured following the addition of 40 μL detection reagent and incubation for 10 min at room temperature using a BioTEK-Flx800 plate reader at 528±20 nm. Determine the amount of GTP consumed in a biochemical reaction by referencing a standard curve that relates light units (RLU) indicative of product formation to GTP concentration. Using this data, calculate the rate of the enzymatic reaction (velocity, v) by applying the Michaelis-Menten equation: v = Vmax/(1+(Km/[S])). To facilitate the determination of *Vmax* and *Km*, construct a Lineweaver-Burk plot, which linearizes the relationship by graphing the reciprocal of the velocity (1/*v*) against the reciprocal of the substrate concentration (1/[*S*]). The calculations of initial rates were performed at 1.08 min in the linear reaction phase of catalysis. Analysis of data was performed by Excel and GraphPad Prism^®^ 9.0.

### BODIPY-FL-GTP, BODIPY-FL-GDP binding, and nucleotide exchange

The nucleotide-binding assay utilized BODIPY-FL-GTP and BODIPY-FL-GDP, from Invitrogen™, each supplied as a mixture of two isomers with the fluorophore attached at either the 2′ or 3′ position on the ribose ring. BODIPY-FL-GTP could be hydrolyzed to BODIPY-FL-GDP. To form the GPCR-G protein complex, a solution containing 2 μM G protein and 100 μM receptor was incubated for 30 minutes at 22°C. The fully activated state of the WT* receptor was achieved by supplementing the mixture with 10 mM NECA. Assays were conducted at 22°C using 96-well half-area microtiter plates in a BioTek plate reader, with excitation at 475 nm and emission measured at 528 nm. The assay buffer comprised 20 mM HEPES (pH 7.4), 1 mM EDTA, and 10 mM MgCl2, supplemented with 0.01% LMNG-3 for protein stability. Initial kinetic data were acquired for 100 nM BODIPY-FL-GTP/BODIPY-FL-GDP in the absence of G protein for 70 seconds to establish a baseline fluorescence intensity. Subsequently, 200 nM heterotrimer G proteins, with or without the receptor, were added, and mixing was rapidly performed in the fluorescence cuvette. Data collection proceeded uninterrupted, and resulting kinetics spectra were plotted and fitted to a one-phase association function using GraphPad Prism 9.0. For the GTP-GDP exchange assay, the GPCR-G protein complex was formed as described, followed by incubation with 100 nM BODIPY-FL-GDP for 2 hours. Baseline fluorescence intensity was measured for 70 seconds using the plate reader, after which 1 μM GTP (in a concentration of 10×GDP), procured from Invitrogen™, was added to facilitate the exchange of BODIPY-FL-GDP to GTP. All experiments were repeated three times, and resulting kinetics spectra were analyzed using GraphPad Prism 9.0.

### Gaussian accelerated molecular dynamics (GaMD)

GaMD is an enhanced sampling method that works by adding a harmonic boost potential to reduce the system energy barriers ^39,40^. When the system potential 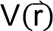 is lower than a reference energy E, the modified potential 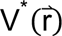 of the system is calculated as:

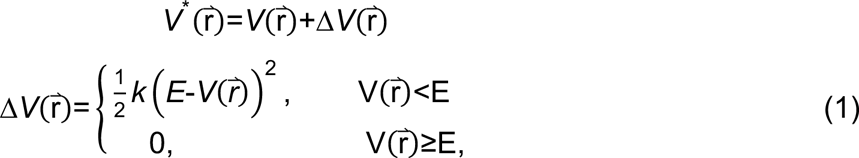

where *k* is the harmonic force constant. The two adjustable parameters E and k are automatically determined on three enhanced sampling principles. First, for any two arbitrary potential values 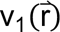 and 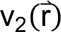 found on the original energy surface, if 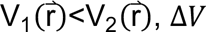, Δ*V* should be a monotonic function that does not change the relative order of the biased potential values; i.e., 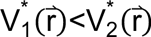. Second, if 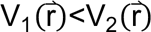, the potential difference observed on the smoothened energy surface should be smaller than that of the original; i.e., 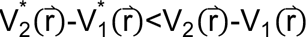. By combining the first two criteria and plugging in the formula of 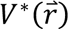 and Δ*V*, we obtain

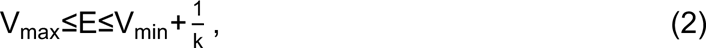

Where V_min_ and V_max_ are the system minimum and maximum potential energies. To ensure that Eq. 2 is valid, *k* has to satisfy: k≤1/(V_max_-V_min_). Let us define: k=k_0_·1/(V_max_-V_min_), then 0<k_0_≤1. Third, the standard deviation (SD) of Δ*V* needs to be small enough (i.e. narrow distribution) to ensure accurate reweighting using cumulant expansion to the second order: σ_ΔV_=k+E-V_avg_,σ_V_≤σ_0_ , where V_avg_ and σ_V_ are the average and SD of ΔVwith σ_0_ as a user-specified upper limit (e.g., 10k_B_T) for accurate reweighting. When E is set to the lower bound E=V_max_ according to Eq. 2, k_0_ can be calculated as

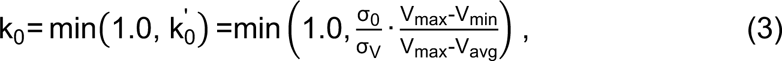

Alternatively, when the threshold energy E is set to its upper bound E=V_min_+1/k, k_0_ is set to:

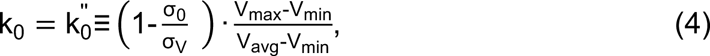

If 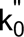 is calculated between 0 and 1. Otherwise, k is calculated using Eq. 3.

### System setup and simulation analysis

The cryo-EM structure of wild-type NECA-bound A2AR bound by mini-Gαs (PDB ID: 6GDG^16^) was used for setting up simulation systems of the wild-type NECA-bound cWT*-A2AR-mini-Gαsβγ, *apo* cR291A-mini-Gαsβγ, *apo* cWT*-A2AR-Gαsβγ and *apo* cR291A-Gαsβγ. The missing residues in the extracellular loop 2 (ECl2) and intracellular loop 3 (ICL3) of the receptor were modelled with Swiss-Modeller^41^. For the *apo* cR291A-mini-Gαsβγ, the simulation structure was generated by mutanting the correspoding R291A and V229C in the wild-type system and deleting the agonist NECA. To build the wild-type *apo* A2AR bound by the full-length of Gs protein (*apo* cWT*-A2AR-Gαsβγ), the Swiss-Modeller^41^ was used to build the Gs protein with a geometry and orientation similar to the Gs protein in the fully active state of the β2AR-Gs complex (PDB ID: 3SN6^1^). The coordinates of the GDP and Mg^2+^ were obtained by aligning the Ras domain of the crystal structure of Gs-bound GDP (PDB ID: 6AU6^42^) to the modelled full-length Gs protein bound by the A2AR.The simulation structure of the *apo* cR291A-Gαsβγ was generated by substituting residues Arg291 and Val232 in the wild-type system (*apo* cWT*-A2AR-Gαsβγ) with Ala and Cys, respectively.

VMD was used to insert the NECA-bound cWT*-A2AR-mini-Gαsβγ, *apo* cR291A-mini-Gαsβγ, *apo* cWT*-A2AR-Gαsβγ, and *apo* cR291A-Gαsβγ complex into POPC (palmitoyl-2-oleoyl-sn-glycero-3-phosphocholine) lipids to prepare simulation systems. In each simulation system, the protein and lipid bilayer were solvated with TIP3P water molecules in a box of 11.2 nm x 13.1 nm x 14.6 nm with the periodic boundary condition. The system charge was neutralized with 150 mM NaCl. The CHARMM36m parameter set^43–45^ was used for the proteins and lipids, and Guanosine diphosphate (GDP). Force field parameters of the NECA agonist were obtained from the ParamChem web server^46^. The four simulation systems were first energy minimized for 5,000 steps with constraints on the heavy atoms of the proteins and phosphor atom of the lipids. The hydrogen-heavy atom bonds were constrained using the SHAKE algorithm and the simulation time step was set to 2.0 fs. The particle mesh Ewald (PME) method^47^ was employed to compute the long-range electrostatic interactions and a cutoff value of 9.0 Å was applied to treat the non-bonded atomic interactions. The temperature was controlled using the Langevin thermostat with a collision frequency of 1.0 ps^-1^. The system was then equilibrate using the constant number, volume, and temperature (NVT) ensemble at 310K for 250 ps and under the constant number, pressure, and temperature (NPT) ensemble at 310 K and 1 bar for another 1 ns with constraints on the heavy atoms of the protein, followed by 10 ns short conventional MD (cMD) without any constraint.

The GaMD module implemented in the GPU version of AMBER22^48–50^ was then applied to perform the simulations of NECA-bound cWT*-A2AR-mini-Gαsβγ, *apo* cR291A-mini-Gαsβγ, *apo* cWT*-A2AR-Gαsβγ, and *apo* cR291A-Gαsβγ. GaMD simulations included a 8-ns short cMD run used to collect the potential statistics for calculating GaMD acceleration parameters, an 56-ns GaMD equilibration after adding the boost potential, and finally three independent 2,000-ns GaMD production simulations with randomized initial atomic velocities for the systems of the NECA-bound cWT*-A2AR-mini-Gαsβγ and *apo* cR291A-A2AR-mini-Gαsβγ complex, and 3,000-ns GaMD production simulations for the systems of *apo* cWT*-A2AR-Gαsβγ and *apo* cR291A-Gαsβγ with randomized initial atomic velocities. The average and SD of the system potential energies were calculated every 800,000 steps (1.6 ns). All GaMD simulations were performed at the “dual-boost” level by setting the reference energy to the lower bound. One boost potential was applied to the dihedral energetic term and the other to the total potential energetic term. The upper limit of the boost potential SD, σ0 was set to 6.0 kcal/mol for both the dihedral and the total potential energetic terms.

For each simulation system, all three GaMD production trajectories were combined together for analysis with CPPTRAJ^51^. The distance between the NPxxY motif of the receptor and the last five residues of Gαs α5 helix (A2AAR:NPxxY-Gαs:α5 distance), the distance between the receptor TM3 and TM6 intracellular ends (measured by the distance between the Cα atoms of receptor residues Arg102^3.50^ and Glu228^6.30^), the root-mean-square derivation of GDP (GDP RMSD) relative to the simulation starting structure were selected as reaction coordinates. The angle between the Ras domain and α helical domain (AHD) was used as another reaction coordinate to indicate their relative orientation, which was defined by the two vectors of Gαs AHD and Gαs Ras domain. Vector 1 went through the Gαs AHD and A161 centers and vector 2 went through Gαs Ras domain and E299 centers. The PyReweighting^52^ toolkit was applied to reweight GaMD simulations to recover the original free energy profiles of the simulation systems. 2D free energy profiles were computed using the combined trajectories from all the three independent GaMD simulations for each system with the A2AR:NPxxY-Gαs:α5 distance, TM3-TM6 distance, GDP RMSD and the angle between the Ras domain and AHD as reaction coordinates. A bin size of 1.0 Å was used for the A2AR:NPxxY-Gαs:α5 distance, TM3-TM6 distance and GDP RMSD. A bin size of 6.0° was used for the angle between the Ras domain and AHD as reaction coordinate. The cutoff was set to 500 frames for 2D free energy calculations.

### Preparation of the A2AR-mini-GSβ1γ2-Nb35 complex

A2AR, mini-GS-β1γ2 and Nb35 were mixed in a molar ratio of 1:2:4, to yield a final complex concentration of 1 mg/mL. To this mixture, 0.1 U of apyrase was added, followed by an overnight incubation at 4°C. The mixture was then concentrated with a 100 kDa MWCO Amicon filter and injected onto a Superdex200 Increase 10/300 GL gel filtration column equilibrated with buffer (50 mM HEPES, 100 mM NaCl, 0.002% LMNG-3, 0.0002% CHS (w/v)). Monodisperse fractions were concentrated with a 100 kDa MWCO Amicon filter immediately prior to cryo-EM grid preparation.

Negative staining of the complex was performed with 0.75% uranium formate, following an established protocol^53^.Grids were examined using an FEI T12 microscope operated at 120 kV, and images were recorded using a 4K x 4K charge-coupled device (CCD) camera (UltraScan 4000, Gatan).

### Cryo-EM sample preparation and data acquisition

Freshly prepared A2AR-mini-GS-β1γ2-Nb35 complex at a final concentration of 1 mg/mL, was applied to glow-discharged gold grids coated with either holey carbon film (Quantifoil, 300 mesh 1.2/1.3, Au) or holey gold film (UltrAuFoil, 300, mesh 1.2/1.3). These grids were then plunge-frozen using a Vitrobot Mark IV with a blotting time of 4 s and blotting force of 0, at 4 °C and 100% humidity. Grids were subsequently examined and screened using an FEI Tecnai Arctica operated at 200 kV and equipped with an XFEG and a Gatan K3 camera. Cryo-EM data collection was performed on a Titan Krios at the UCSF Cryo-EM Center for Structural Biology, operated at an acceleration voltage of 300 kV, equipped with an XFEG, a BioQuantum energy filter (slit width set to 20 eV) and a K3 camera (Gatan).

All cryo-EM datasets were collected using SerialEM^54^. A multishot collection (3×3 arrays) was employed, incorporating beam-tilt compensation and maximum image shift is 3.5 micros. All images were acquired with a nominal magnification of 105 K, resulting in a super-resolution pixel size of 0.4175 Å (physical pixel size of 0.835 Å). Defocus range was set from -0.8 μm to -1.8 μm. A total of 8,805 images were collected, each was dose-fractionated into 80 movie frames with a total exposure time of 2.024 s, resulting in a total fluence of approximately 47.7 electrons per Å^2^. Data collection statistics are shown in Supplementary Table 1.

### Image process

A total of 8,805 movie stacks were motion-corrected, dose-weighted, and binned by Fourier cropping to the physical pixel size of 0.835 Å on-the-fly using MotionCor2^55^. Motion-corrected, does-weighted sums were used for contrast transfer function (CTF) determination and resolution estimation in cryoSPARC^56^. 424,536 particles were picked from randomly selected 500 micrographs using cryoSPARC blob picker with a diameter of 80-150 Å. These particles were subjected to ab-initio reconstruction and multi-round heterogenous refinement. One class with distinct features of A2AR bound to mini-Gαsβ1γ2-Nb35 was identified, from which 16 different projection images were created for template picking, yielding 7,282,317 particles from all micrographs.

After removing junk particles by extensive 2D classifications, 2,328,810 particles were selected for ab-initio reconstruction and multi-round heterogenous refinement. One distinct class of 307,568 particles was identified for further non-uniform refinement of the A2AR-mini-Gαsβ1γ2-Nb35 complex, yielding a reconstruction with a global resolution of 3.04Å. This particle stack was then exported to RELION^57^ for multiple rounds of 3D refinement (initial low-pass filter: 10 Å; mask diameter: 360 Å; reference mask: no) to produce a new reconstruction, from which a mask without the detergent micelle was generated by using the segment map function in Chimera^58^ (initial threshold: 0.0001; extend_inimask: 4; width_soft_edge: 4). Particle subtraction function in RELION was applied by using this mask to generate a micelle removed particle stack.

Next, we applied 3D classification to the micelle subtracted particles with a reference model without micelle (reference mask: generated last step; initial low-pass filter: 10 Å; mask diameter: 260 Å; regularisation parameter T: 3; number of iterations: 50; number of classes: 5; perform local angular searches: no). A new reconstruction of the entire A2AR-mini-GS-β1γ2-Nb35 complex from 71,547 particles was identified for further refinement in RELION (initial low-pass filter: 3.5 Å; mask diameter: 260 Å; angular sampling interval: 1.8°; local search from auto-sampling: 0.9°). A final 3D reconstruction was calculated in cisTEM^59^ with default settings and without further refinement. The numeric resolution was determined from Fourier Shell Correction (FSC) using criterion of FSC = 0.143^60^. The final map was sharpened by DeepEMhancer^61^ and used for model building and figure generation.

### Model building

For model building, the initial model was generated by fitting the existing coordinates of the activated state of A2AR-mini-GS-β1γ2-Nb35 complex (PDB: 6GDG) into our cryo-EM density maps using ChimeraX^62^. Discrepancies between the initial model and the density maps were then manually built and refined ISOLDE^63^ and coot^64^. Subsequent refinements were performed in Phenix^65^ with secondary structure constraints. The model was validated by wwPDB validation server^66^ and no major issues was reported. A summary of the parameters used in data collection and model building is provided in Supplementary Table 1.

### Data representation

All statistical tests such as GTP hydrolysis assessments were conducted using GraphPad Prism 9.0. The central point of all data points gives the mean value with s.d. for all data unless otherwise specified. Visualization of the atomic models (figures and videos) were made using UCSF ChimeraX and PyMoL (The PyMOL Molecular Graphics System, Version 2.0 Schrödinger, LLC.)

## Data Availability

The atomic coordinates of intermediate A2AR-mini-Gαsβγ has been deposited in the Protein Data Bank with ID: 8VM3.

## Acknowledgements

We thank Drs. Christopher G. Tate and Javier García Nafría for providing the plasmid of mini-G399. We also thank Hiran Malinda Lamabadu Warnakulasurya and Natasha Jaiswal in assistance of GTP hydrolysis measurements, Dr. Hao Wu for technical assistance in cryo-EM data acquisition and advice on data processing and model building, Wooyoung Choi, Will Arnold and Adamo Mancino for their advice and feedback on figures, and to the members of the Cheng laboratory for helpful discussions. The cryo-EM facility at UCSF is managed by David Bulkley and Glenn Gilbert. Some of this work was performed at the Janelia cryo-EM facility, supported by Howard Hughes Medical Institute (HHMI). Computation at the Cheng laboratory is supported by Chengmin Li and Matt Harrington. This work is partly supported by grant from National Institute of Health (5R01GM132572(Y. M.), 5R35GM140847(Y.C.), and 1R01GM149659 (L. Y.)). Y.C. is also an investigator of Howard Hughes Medical Institute.

## Author contributions

L. Y. conceptualized the study. M.B prepared and purified complex for cryo-EM and performed all structural analysis. X. W. expressed and purified the receptors and different component proteins. X. W. also performed NMR spectral acquisition, data process and spectral analyses in addition to biochemical assessments. J. W. performed the MD simulation. J. X prepared G proteins. W.S prepared part of the receptors and conducted biochemical assays. V.A.A was involved in MD simulation. Y.M. supervised the MD simulation. Y.C supervised the structural analysis. L. Y. supervised receptor and G protein purification, NMR acquisition, and spectral analysis, and biochemical assays. All authors participated in manuscript writing and revisions.

## Competing interests

The authors declare no competing financial interests.

## Extended Data Figures

**Extended Data Fig. 1:**
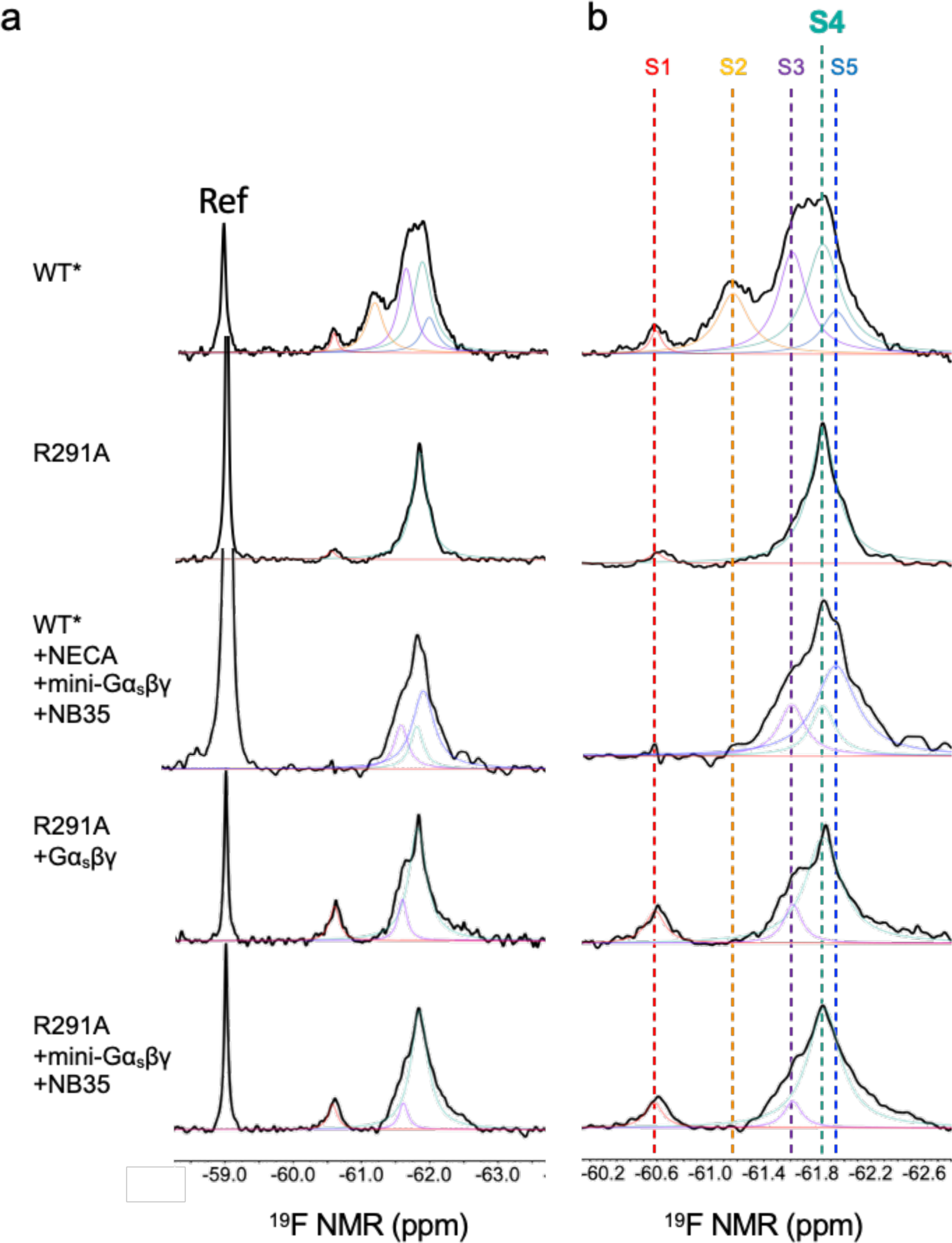
Conformational profiles of the mutant R291A and its complexes probed by ^19^F-qNMR. **(A)** and **(B),** Deconvoluted conformational profiles probed by ^19^F-qNMR as a function of mini-Gα_s_βγ and Gα_s_βγ (S4-Gαβγ), in reference to the WT*-mini-Gα_s_βγ+NB35 (S5-Gα_s_βγ), representing the conformational profile for the structure PDB ID: 6GDG.

**Extended Data Fig. 2:**
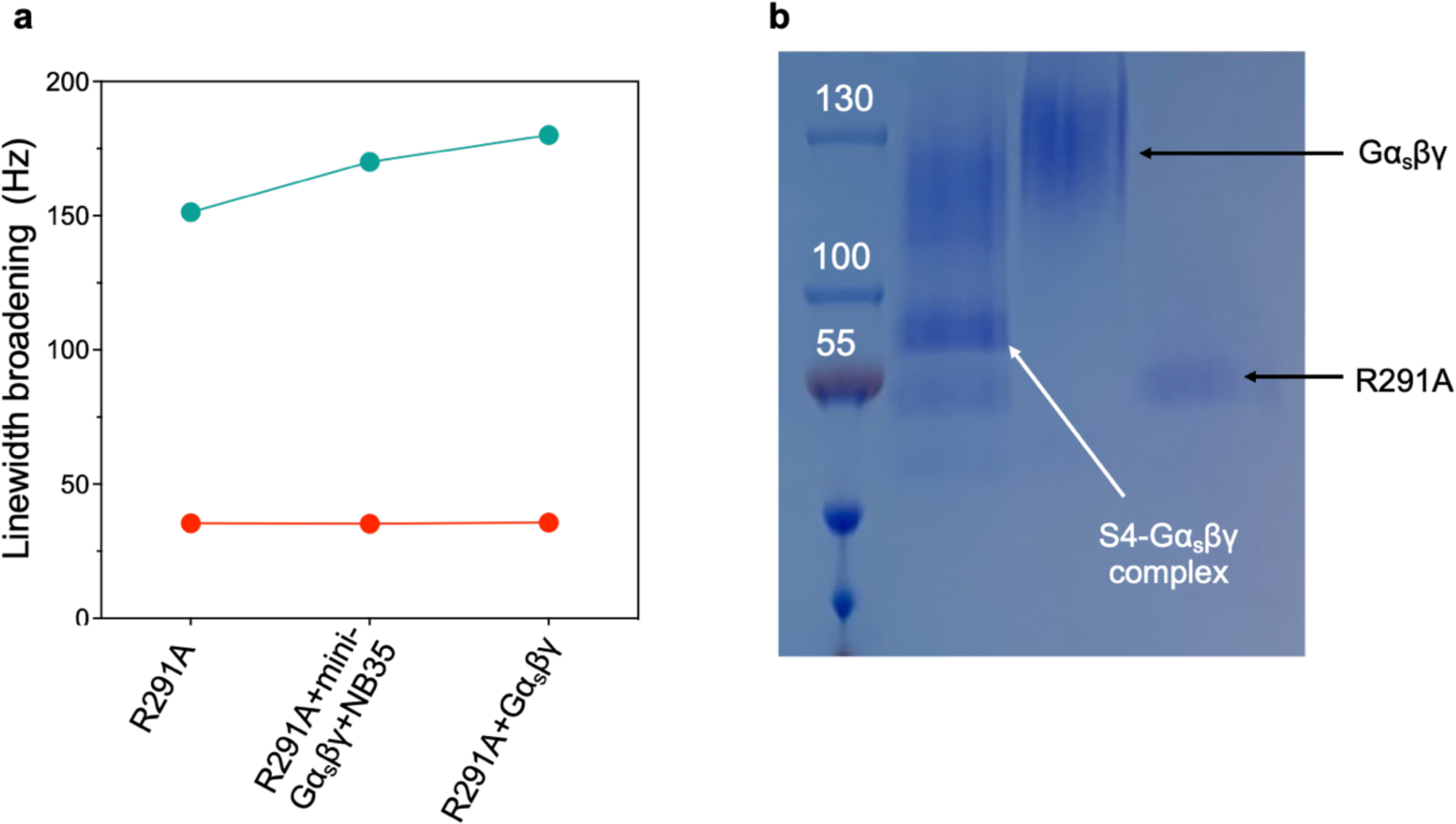
Both linewidth broadening of ^19^F-qNMR resonance for S4 intermediate and native-PAGE indicated the complex formation between the S4 and G protein. **a,** Linewidths of S4 resonance for R291A, R291A-Gα_s_βγ, and R291A-mini-Gα_s_βγ-NB35. **b,** Native-PAGE for R291A and Gα_s_βγ interaction.

**Extended Data Fig. 3:**
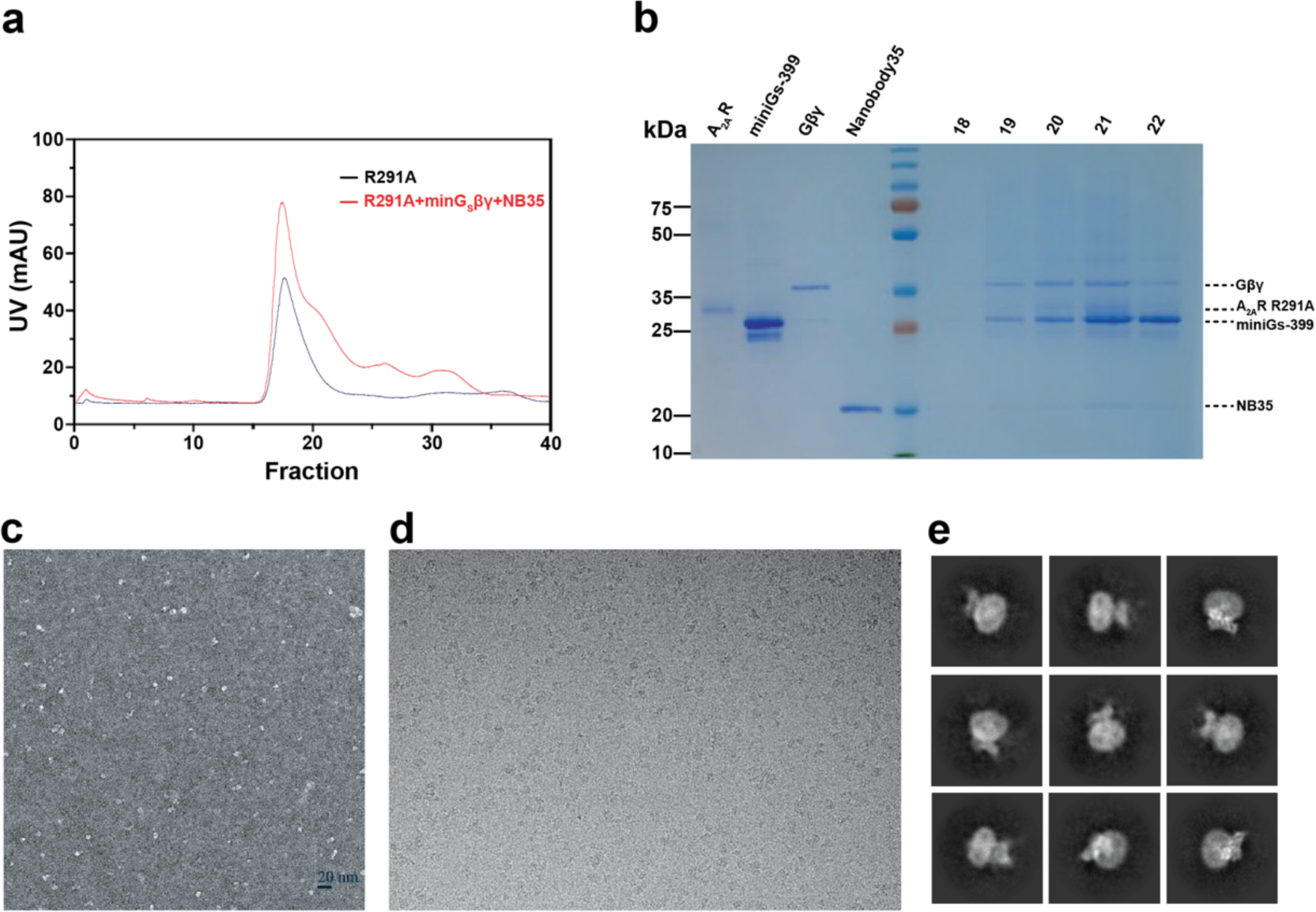
Preparation of A_2A_R sample and cryo-EM screening. **a,** Size exclusion chromatography of purified A2AR-R291A with and without mini-G_s_βγ+NB35. **b,** SDS-PAGE of the indicated fraction, corresponding to R291A+mini-G_s_βγ+NB35. **c**, Negative staining of the R291A+ mini-G_s_βγ+NB35 complex. **d**, Representative cryo-EM micrograph of the complex. **e**, Representative 2D average image of the complex.

**Extended Data Fig. 4:**
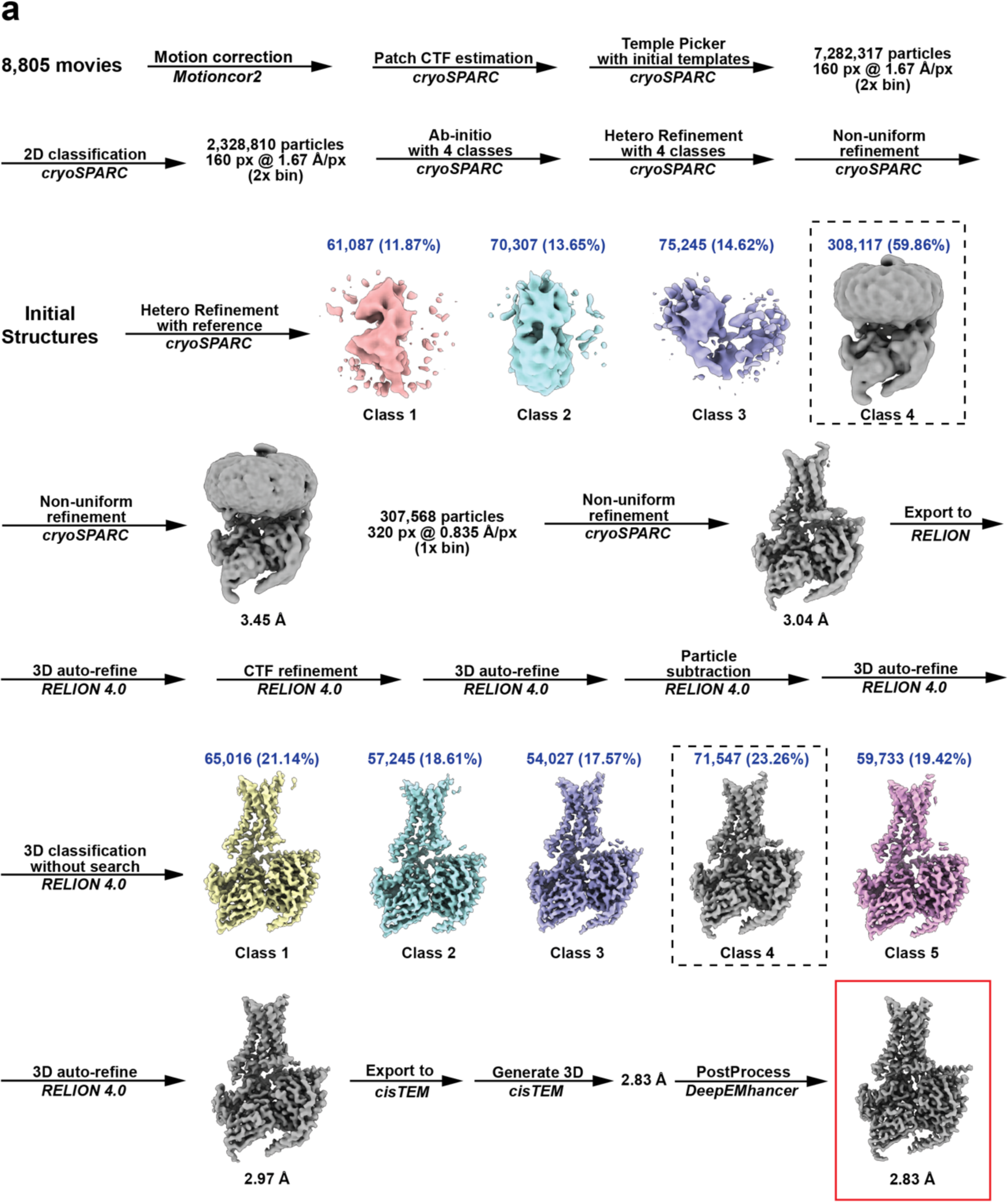
Cryo-EM data and image processing flow. Schematic flow-chat representing the image processing approach for A_2A_R-R291A. Thumbnail images of each 3D class or refinement are shown along with global GS-FSC resolution in black, particles counts in blue, and dashed black boxes to indicate selected 3D classes. Cryo-EM map (red box) and atomic model are used in main figures to present A_2A_R-R291A features.

**Extended Data Fig. 5:**
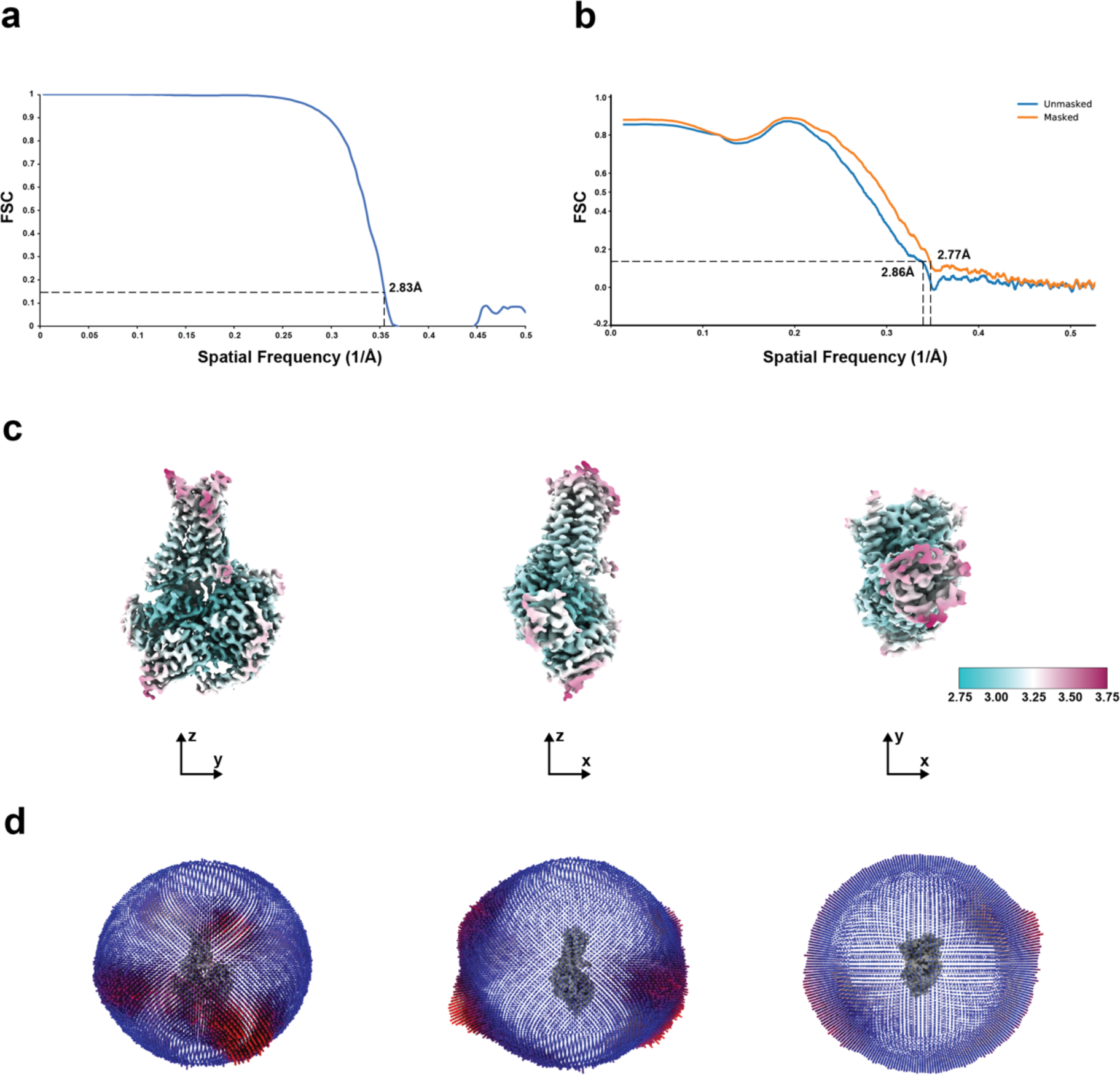
Resolution estimation and atomic model quality. **a,** Fourier shell correlation curve with resolution determined by the ‘gold-standard’ FSC. **b,** Model-to-map FSC plots calculated by PHENIX between the map and the model. **c,** Cryo-EM map colored by local resolution estimates using RELION with resolution scale bar (Å). **d**, Histogram representation of the Euler angle distribution from the final particles used in the reconstruction.

**Extended Data Fig. 6:**
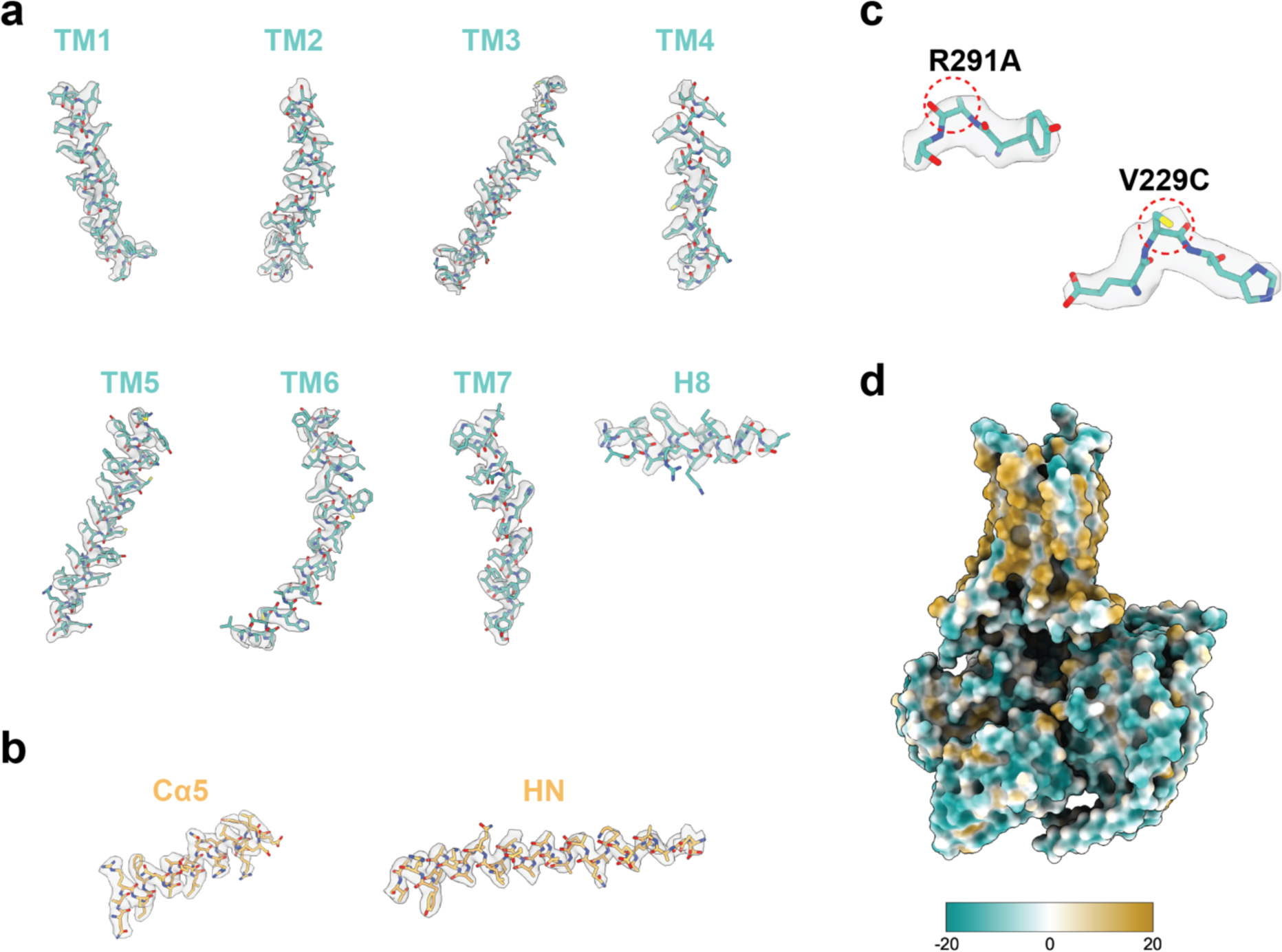
Representative cryo-EM densities from selected structural features of A_2A_R. **a**, Densities of all transmembrane helices. **b**, Representative densities of mini-G. **c,** Cryo-EM densities of V229C and R291A. **d,** Representation of surface hydrophobicity.

**Extended Data Fig. 7:**
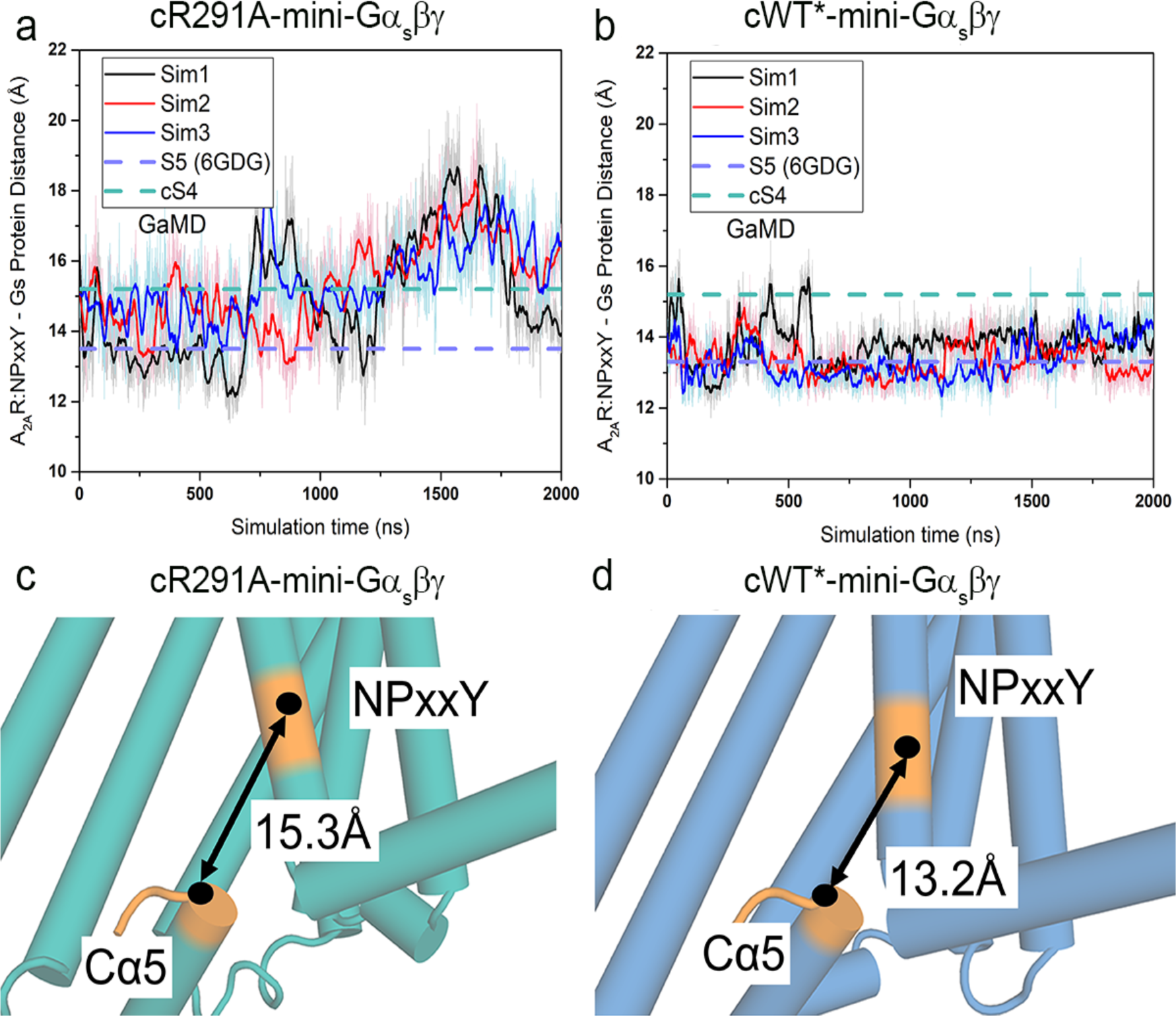
GaMD simulations for cR291A-mini-Gα_s_βγ and cWT*-mini-Gα_s_βγ. **a-b**, Time course of the center-of-mass (COM) distance between the receptor NPxxY motif in TM7 and the Cα5 helix (A_2A_R:NPxxY-Gαs:Cα5 distance) calculated from the GaMD simulations of cR291A-mini-Gα_s_βγ (**a**) and cWT*-mini-Gα_s_βγ (**b**) complex systems. **c-d**, Representative low-energy conformations of the cR291A-mini-Gα_s_βγ (**c**) and **(d)** cWT*-mini-Gα_s_βγ complex identified from the GaMD simulations. The receptor NPxxY motif and the last five residues of Gα5 helix were colored in orange.

**Extended Data Fig. 8:**
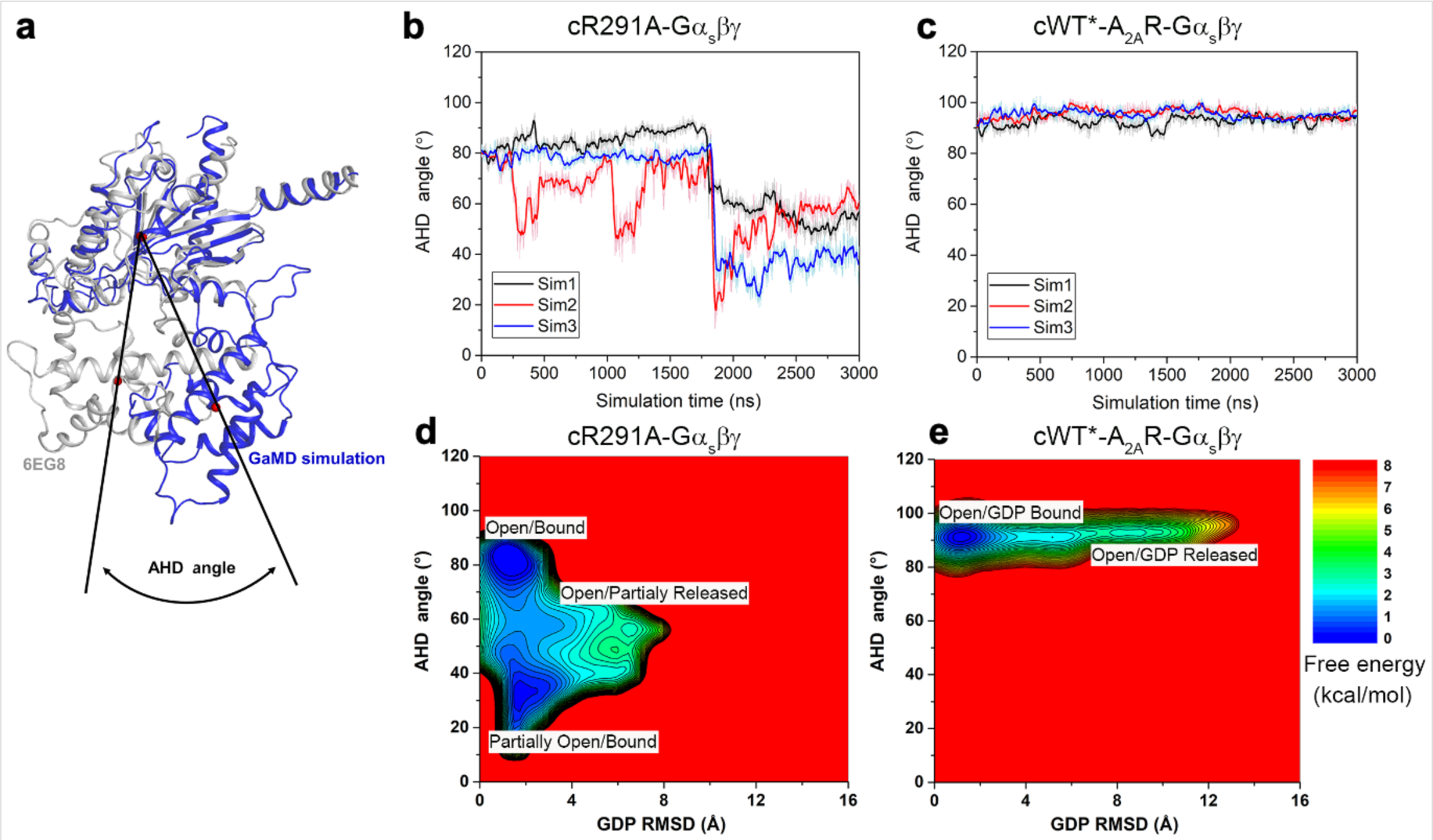
The behaviors of AHD domain in cS4 and cS5 mediated G protein. **a,** The AHD angle is illustrated by comparing the GaMD simulation conformation with the inactive conformation of Gα_s_ (PDB: 6EG8); Vector 1 goes through centers of the Gα_s_ AHD and Ras-like domain of the inactive conformation (6EG8). Vector 2 goes through the centers of the Gα_s_ AHD and Ras-like domain of the GaMD simulation conformations after aligning the Ras-like domain to the inactive conformation of Gα_s_. **b**-**c**, The time course of the AHD angle calculated from the GaMD simulations for the *apo* cR291A-Gα_s_βγ (**b**) and *apo* cWT*-A_2A_R-Gα_s_βγ (**c**) complexes. **d**-**e**, 2D free energy profiles of the *apo* cR291A-Gα_s_βγ (**d**) and *apo* cWT*-Gα_s_βγ (**e**) complexes calculated from GaMD simulations regarding the GDP RMSD and the AHD angle.

## Notes

### Competing Interest Statement

The authors have declared no competing interest.

### Summary of Updates

slightly change abstract and the flow of the manuscript

## References

1 Rasmussen, S. G. et al. Crystal structure of the beta2 adrenergic receptor-Gs protein complex. Nature 477, 549–555 (2011). 10.1038/nature10361

2 Ye, L., Wang, X., McFarland, A. & Madsen, J. J. (19)F NMR: A promising tool for dynamic conformational studies of G protein-coupled receptors. Structure 30, 1372–1384 (2022). 10.1016/j.str.2022.08.007

3 Zhang, Y. et al. Cryo-EM structure of the activated GLP-1 receptor in complex with a G protein. Nature 546, 248–253 (2017). 10.1038/nature22394

4 Hilger, D. The role of structural dynamics in GPCR-mediated signaling. FEBS J 288, 2461–2489 (2021). 10.1111/febs.15841

5 Wang, X., McFarland, A., Madsen, J. J., Aalo, E. & Ye, L. The potential of (19)F NMR application in GPCR biased drug discovery. Trends Pharmacol Sci 42, 19–30 (2021). 10.1016/j.tips.2020.11.001

6 Heo, L. & Feig, M. Multi-state modeling of G-protein coupled receptors at experimental accuracy. Proteins (2022). 10.1002/prot.26382

7 Ye, L., Van Eps, N., Zimmer, M., Ernst, O. P. & Prosser, R. S. Activation of the A2A adenosine G-protein-coupled receptor by conformational selection. Nature 533, 265–268 (2016). 10.1038/nature17668

8 Ye, L., Orazietti, A. P., Pandey, A. & Prosser, R. S. High-efficiency expression of yeast-derived G-protein coupled receptors and (19)F labeling for dynamical studies. Methods Mol Biol 1688, 407–421 (2018). 10.1007/978-1-4939-7386-6_19

9 Perez, D. M. & Karnik, S. S. Multiple signaling states of G-protein-coupled receptors. Pharmacological reviews 57, 147–161 (2005). 10.1124/pr.57.2.2

10 Marti-Solano, M., Schmidt, D., Kolb, P. & Selent, J. Drugging specific conformational states of GPCRs: challenges and opportunities for computational chemistry. Drug discovery today 21, 625–631 (2016). 10.1016/j.drudis.2016.01.009

11 Dong, S. S., Goddard, W. A., 3rd & Abrol, R. Identifying multiple active conformations in the G protein-coupled receptor activation landscape using computational methods. Methods in cell biology 142, 173–186 (2017). 10.1016/bs.mcb.2017.07.009

12 Madurga, S., Nedyalkova, M., Mas, F. & Garces, J. L. Ionization and conformational equilibria of citric acid: delocalized proton binding in solution. The journal of physical chemistry. A 121, 5894–5906 (2017). 10.1021/acs.jpca.7b05089

13 Wang, X., Neale, C., Kim, S. K., Goddard, W. A. & Ye, L. Intermediate-state-trapped mutants pinpoint G protein-coupled receptor conformational allostery. Nat Commun 14, 1325 (2023). 10.1038/s41467-023-36971-6

14 Dore, A. S. et al. Structure of the adenosine A2A receptor in complex with ZM241385 and the xanthines XAC and caffeine. Structure 19, 1283–1293 (2011). 10.1016/j.str.2011.06.014

15 Jaakola, V. P. et al. The 2.6 angstrom crystal structure of a human A2A adenosine receptor bound to an antagonist. Science 322, 1211–1217 (2008). 10.1126/science.1164772

16 Garcia-Nafria, J., Lee, Y., Bai, X., Carpenter, B. & Tate, C. G. Cryo-EM structure of the adenosine A(2A) receptor coupled to an engineered heterotrimeric G protein. Elife 7 (2018). 10.7554/eLife.35946

17 Kenakin, T. New concepts in pharmacological efficacy at 7TM receptors: IUPHAR review 2. Br J Pharmacol 168, 554–575 (2013). 10.1111/j.1476-5381.2012.02223.x

18 Mahoney, J. P. & Sunahara, R. K. Mechanistic insights into GPCR-G protein interactions. Curr Opin Struct Biol 41, 247–254 (2016). 10.1016/j.sbi.2016.11.005

19 Huang, S. K. et al. Allosteric modulation of the adenosine A(2A) receptor by cholesterol. Elife 11 (2022). 10.7554/eLife.73901

20 Huang, S. K. et al. Delineating the conformational landscape of the adenosine A(2A) receptor during G protein coupling. Cell 184, 1884–1894 e1814 (2021). 10.1016/j.cell.2021.02.041

21 Du, Y. et al. Assembly of a GPCR-G protein complex. Cell 177, 1232–1242 e1211 (2019). 10.1016/j.cell.2019.04.022

22 Bai, C. et al. Exploring the Activation Process of the beta2AR-Gs Complex. J Am Chem Soc 143, 11044–11051 (2021). 10.1021/jacs.1c03696

23 Wacker, D. et al. Structural features for functional selectivity at serotonin receptors. Science 340, 615–619 (2013). 10.1126/science.1232808

24 Zondlo, N. J. Aromatic-proline interactions: electronically tunable CH/pi interactions. Acc Chem Res 46, 1039–1049 (2013). 10.1021/ar300087y

25 Shi, L. et al. Beta2 adrenergic receptor activation. Modulation of the proline kink in transmembrane 6 by a rotamer toggle switch. J Biol Chem 277, 40989–40996 (2002). 10.1074/jbc.M206801200

26 Ragnarsson, L., Andersson, A., Thomas, W. G. & Lewis, R. J. Mutations in the NPxxY motif stabilize pharmacologically distinct conformational states of the alpha(1B)- and beta(2)-adrenoceptors. Sci Signal 12 (2019). 10.1126/scisignal.aas9485

27 Cherezov, V. et al. High-resolution crystal structure of an engineered human beta2- adrenergic G protein-coupled receptor. Science 318, 1258–1265 (2007). 10.1126/science.1150577

28 Westfield, G. H. et al. Structural flexibility of the G alpha s alpha-helical domain in the beta2-adrenoceptor Gs complex. Proc Natl Acad Sci U S A 108, 16086–16091 (2011). 10.1073/pnas.1113645108

29 Duan, J. et al. Structures of full-length glycoprotein hormone receptor signalling complexes. Nature 598, 688–692 (2021). 10.1038/s41586-021-03924-2

30. J., G. A. An introduction of Hydrogen bonding. Oxford Unviersity Press (1998).

31 Papasergi-Scott, M. M. et al. Time-resolved cryo-EM of G-protein activation by a GPCR. Nature (2024). 10.1038/s41586-024-07153-1

32 Flock, T. et al. Universal allosteric mechanism for Galpha activation by GPCRs. Nature 524, 173–179 (2015). 10.1038/nature14663

33 Papasergi-Scott, M. M. et al. Time-resolved cryo-EM of G protein activation by a GPCR. bioRxiv (2023). 10.1101/2023.03.20.533387

34 Dror, R. O. et al. SIGNAL TRANSDUCTION. Structural basis for nucleotide exchange in heterotrimeric G proteins. Science 348, 1361–1365 (2015). 10.1126/science.aaa5264

35 Zhao, W., Wang, X. & Ye, L. Expression and purification of yeast-derived GPCR, Gα and Gβγ subunits for structural and dynamic studies. Bio Protoc 11, e3919 (2021). 10.21769/BioProtoc.3919

36 Carpenter, B. & Tate, C. G. Engineering a minimal G protein to facilitate crystallisation of G protein-coupled receptors in their active conformation. Protein Eng Des Sel 29, 583–594 (2016). 10.1093/protein/gzw049

37 Carpenter, B. & Tate, C. G. Expression and purification of mini G proteins from Escherichia coli. Bio-protocol 7, e2235–e2235 (2017).

38 Mondal, S., Hsiao, K. & Goueli, S. A. A homogenous bioluminescent system for measuring GTPase, GTPase activating protein, and guanine nucleotide exchange factor activities. Assay Drug Dev Technol 13, 444–455 (2015). 10.1089/adt.2015.643

39 Wang, J. N. et al. Gaussian accelerated molecular dynamics: Principles and applications. Wires Comput Mol Sci 11, e1521 (2021). 10.1002/wcms.1521

40 Miao, Y., Feher, V. A. & McCammon, J. A. Gaussian Accelerated Molecular Dynamics: Unconstrained Enhanced Sampling and Free Energy Calculation. Journal of Chemical Theory and Computation 11, 3584–3595 (2015). 10.1021/acs.jctc.5b00436

41 Waterhouse, A. et al. SWISS-MODEL: homology modelling of protein structures and complexes. Nucleic Acids Res 46, W296–W303 (2018). 10.1093/nar/gky427

42 Hu, Q. & Shokat, K. M. Disease-Causing Mutations in the G Protein Gαs Subvert the Roles of GDP and GTP. Cell 173, 1254–1264.e1211 (2018). 10.1016/j.cell.2018.03.018

43 Vanommeslaeghe, K. & MacKerell, A. D. CHARMM additive and polarizable force fields for biophysics and computer-aided drug design. Biochimica et Biophysica Acta (BBA) - General Subjects 1850, 861–871 (2015). 10.1016/j.bbagen.2014.08.004

44 Huang, J. et al. CHARMM36m: an improved force field for folded and intrinsically disordered proteins. Nature Methods 14, 71 (2016). 10.1038/nmeth.4067

45 Klauda, J. B. et al. Update of the CHARMM All-Atom Additive Force Field for Lipids: Validation on Six Lipid Types. The Journal of Physical Chemistry B 114, 7830–7843 (2010). 10.1021/jp101759q

46 Vanommeslaeghe, K. & MacKerell, A. D. Automation of the CHARMM general force field (CGenFF) I: Bond perception and atom typing. Journal of Chemical Information and Modeling 52, 3144–3154 (2012). 10.1021/ci300363c

47 Darden, T., York, D. & Pedersen, L. Particle mesh Ewald: An N⋅ log (N) method for Ewald sums in large systems. J Chem Physics 98, 10089 (1993).

48 D.A. Case, D.S. Cerutti, T.E. Cheatham, III, T.A. Darden, R.E. Duke, T.J. Giese, H. Gohlke, A.W. Goetz, D. Greene, N. Homeyer, S. Izadi, A. Kovalenko, T.S. Lee, S. LeGrand, P. Li, C. Lin, J. Liu, T. Luchko, R. Luo, D. Mermelstein, K.M. Merz, G. Monard, H. Nguyen, I. Omelyan, A. Onufriev, F. Pan, R. Qi, D.R. Roe, A. Roitberg, C. Sagui, C.L. Simmerling, W.M. Botello-Smith, J. Swails, R.C. Walker, J. Wang, J. Wang, R.M. Wolf, X. Wu, L. Xiao, D.M. York and P.A. Kollman (2022), AMBER 2022, University of California, San Francisco.

49 Miao, Y., Feher, V. A. & McCammon, J. A. Gaussian Accelerated Molecular Dynamics: Unconstrained Enhanced Sampling and Free Energy Calculation. J Chem Theor Comput 11, 3584–3595 (2015). 10.1021/acs.jctc.5b00436

50 Wang, J. et al. Gaussian accelerated molecular dynamics (GaMD): principles and applications. Wiley Interdiscip Rev Comput Mol Sci 11, e1521 (2021). 10.1002/wcms.1521

51 Roe, D. R. & Cheatham, T. E., 3rd. PTRAJ and CPPTRAJ: software for processing and analysis of molecular dynamics trajectory data. J. Chem. Theory Comput. 9, 3084–3095 (2013). 10.1021/ct400341p

52 Miao, Y. et al. Improved reweighting of accelerated molecular dynamics simulations for free energy calculation. J. Chem. Theory Comput. 10, 2677–2689 (2014). 10.1021/ct500090q

53 Ohi, M., Li, Y., Cheng, Y. & Walz, T. Negative staining and image classification - powerful tools in modern electron microscopy. Biol Proced Online 6, 23–34 (2004). 10.1251/bpo70

54 Mastronarde, D. N. Automated electron microscope tomography using robust prediction of specimen movements. J Struct Biol 152, 36–51 (2005). 10.1016/j.jsb.2005.07.007

55 Zheng, S. Q. et al. MotionCor2: anisotropic correction of beam-induced motion for improved cryo-electron microscopy. Nat Methods 14, 331–332 (2017). 10.1038/nmeth.4193

56 Punjani, A., Rubinstein, J. L., Fleet, D. J. & Brubaker, M. A. cryoSPARC: algorithms for rapid unsupervised cryo-EM structure determination. Nat Methods 14, 290–296 (2017). 10.1038/nmeth.4169

57 Scheres, S. H. RELION: implementation of a Bayesian approach to cryo-EM structure determination. J Struct Biol 180, 519–530 (2012). 10.1016/j.jsb.2012.09.006

58 Pettersen, E. F. et al. UCSF Chimera--a visualization system for exploratory research and analysis. J Comput Chem 25, 1605–1612 (2004). 10.1002/jcc.20084

59 Grant, T., Rohou, A. & Grigorieff, N. cisTEM, user-friendly software for single-particle image processing. Elife 7 (2018). 10.7554/eLife.35383

60 Rosenthal, P. B. & Henderson, R. Optimal determination of particle orientation, absolute hand, and contrast loss in single-particle electron cryomicroscopy. J Mol Biol 333, 721–745 (2003). 10.1016/j.jmb.2003.07.013

61 Sanchez-Garcia, R. et al. DeepEMhancer: a deep learning solution for cryo-EM volume post-processing. Commun Biol 4, 874 (2021). 10.1038/s42003-021-02399-1

62 Goddard, T. D. et al. UCSF ChimeraX: Meeting modern challenges in visualization and analysis. Protein Sci 27, 14–25 (2018). 10.1002/pro.3235

63 Croll, T. I. ISOLDE: a physically realistic environment for model building into low-resolution electron-density maps. Acta Crystallogr D Struct Biol 74, 519–530 (2018). 10.1107/S2059798318002425

64 Emsley, P. & Cowtan, K. Coot: model-building tools for molecular graphics. Acta Crystallogr D Biol Crystallogr 60, 2126–2132 (2004). 10.1107/S0907444904019158

65 Afonine, P. V. et al. Towards automated crystallographic structure refinement with phenix.refine. Acta Crystallogr D Biol Crystallogr 68, 352–367 (2012). 10.1107/S0907444912001308

66 Berman, H., Henrick, K. & Nakamura, H. Announcing the worldwide Protein Data Bank. Nat Struct Biol 10, 980 (2003). 10.1038/nsb1203-980

