## Supplementary Table 1 for "Structure and function of an intermediate GPCR-Gαβγ complex"

**Supplementary Table 1|Cryo-EM data collection, refinement, and validation statistics**

| | $A_{2A}$R-R291A<br>EMDB-43351<br>PDB 8VM3 |
|---|---|
| **Data collection and processing** | |
| Microscope | Titan Krios |
| Voltage (keV) | 300 |
| Camera | Gatan K3 with Gatan Bioquantum energy filter |
| Nominal/Calibrated Magnification | 105,000 |
| Pixel size (Å) | 0.835 |
| Total electron exposure ($e^-$/Å$^2$) | 47.7 |
| Exposure rate ($e^-$/pixel/sec) | 16 |
| Defocus range (μM) | -0.8-1.8 |
| Automation software | SerialEM |
| Energy filter slit width (eV) | 20 |
| Micrographs used (no.) | 8,805 |
| Total extracted/refined particles (no.) | 7,282,317/307,568 |
| **Reconstruction** | |
| Final particles | 71,547 |
| Symmetry | C1 |
| Resolution (global, Å) | 2.83 |
| Resolution (global, Å) | |
| FSC 0.5 (unmasked/masked) | 3.44/3.21 |
| FSC 0.143 (unmasked/masked) | 2.86/2.77 |
| Resolution range (local, Å) | 2.75-3.75 |
| Map sharpening *B* factor (Å$^2$) | -81 |
| **Model composition** | |
| Chains | 5 |
| Atoms | 14828 (Hydrogens: 7193) |
| Protein residues | 1025 |
| Ligands | 0 |
| **Model Refinement** | |
| Refinement package | |
| -real or reciprocal space | Real space |
| -resolution cutoff | 0.143 |
| Model-Map scores | |
| -CC | 0.84 |
| *B* factor (Å$^2$) | |
| Protein (min/max/mean) | 16.69/113.24/54.03 |
| Ligands | N/A |
| R.m.s. deviations from ideal values | |
| Bond lengths (Å) | 0.005 |
| Bond angles (°) | 0.835 |
| **Validation** | |
| MolProbity score | 1.59 |
| CaBLAM outliers (%) | 1.91 |
| Clash score | 5.40 |
| Poor rotamers (%) | 0.13 |
| Ramachandran plot | |
| Favored (%) | 95.74 |
| Allowed (%) | 4.26 |
| Outliers (%) | 0.00 |
